# Advected percolation in the actomyosin cortex drives amoeboid cell motility

**DOI:** 10.1101/2022.07.14.500109

**Authors:** Juan Manuel García-Arcos, Johannes Ziegler, Silvia Grigolon, Loïc Reymond, Gaurav Shajepal, Cédric J. Cattin, Alexis Lomakin, Daniel Müller, Verena Ruprecht, Stefan Wieser, Raphael Voituriez, Matthieu Piel

**Affiliations:** Institut Pierre Gilles de Gennes, PSL Research University. 6 rue Jean Calvin, 75005, Paris, France; Institut Curie, PSL Research University, CNRS, UMR 144, Paris, France; University of Geneva, Department of Biochemistry. Sciences II, Quai Ernest-Ansermet 30, 1205 Geneva, Switzerland; ICFO – Institut de Ciencies Fotoniques, The Barcelona Institute of Science and Technology, 08860 Castelldefels, Spain; IRIS Technology Solutions S.L., Carretera d’Esplugues 39-41, 08940 Cornellà de Llobregat (Barcelona), Spain; Sorbonne Université, CNRS, Institut de Biologie Paris-Seine (IBPS), Laboratoire Jean Perrin (LJP), F-75005, Paris, France; Centre for Genomic Regulation (CRG), The Barcelona Institute of Science and Technology, 08003 Barcelona, Spain; Department of Biosystems Science and Engineering, ETH Zurich, 4058 Basel, Switzerland; F. Hoffmann-La Roche Ltd., Basel, Switzerland; Center for Pathobiochemistry and Genetics, Institute of Medical Chemistry, Medical University of Vienna, Währingerstraße 10, 1090 Vienna, Austria; Center for Pathobiochemistry and Genetics, Institute of Medical Genetics, Medical University of Vienna, Währingerstraße 10, 1090 Vienna, Austria; Universitat Pompeu Fabra (UPF), Barcelona, Spain; Laboratoire de Physique Théorique de la Matière Condensée, CNRS/Sorbonne Université, 4 Place Jussieu, 75005 Paris, France

## Abstract

Spontaneous locomotion is a common feature of most metazoan cells, generally attributed to the fundamental properties of the actomyosin network. This force-producing machinery has been studied down to the most minute molecular details, especially in lamellipodium-driven migration. Nevertheless, how actomyosin networks work inside contraction-driven amoeboid cells still lacks unifying principles. Here, using stable motile blebs as a model amoeboid motile system, we image the dynamics of the actin cortex at the single filament level and reveal the co-existence of three phases of the actin network with distinct rheological properties. Physical modelling shows that these three phases organize spontaneously due to a rigidity percolation transition combined with an active advection of the percolated network. This spontaneous spatial organization of the mechanical properties of the actin network, which we propose to call advected percolation, constitutes a minimal and generic locomotion mechanism. It explains, down to the single actin filament level and up to the scale of the entire cell, how amoeboid cells can propel efficiently through complex 3D environments, a feature shared by immune and cancer cells.

## Introduction

Most animal tissue cells display the intrinsic propensity for spontaneous motility when placed in the appropriate environment, even if they are normally immobile in their tissue of origin (Liu et al., 2015; Ruprecht et al., 2015). The reawakening of the dormant motile capacity of single cells is also thought to be responsible for the dissemination of many different types of cancer cells, for example in the context of the epithelial-to-mesenchymal transition, leading to metastasis (Friedl and Wolf, 2003). This locomotory capacity of single cells has been studied for decades and is generally attributed to the fundamental properties of the actomyosin network, which can spontaneously polarize and produce directed forces (Ierushalmi et al., 2020; Kapustina et al., 2013; Yam et al., 2007).

Experimental as well as theoretical work have converged to define the general properties of actomyosin systems, both *in vitro* and inside cells (Alvarado et al., 2017; Jülicher et al., 2007; Koenderink and Paluch, 2018; Prost et al., 2015). These studies have provided phase diagrams relating the composition of the network (the density of filaments, crosslinkers, and motors) to its large-scale organization and dynamics (Alvarado et al., 2013; Kruse et al., 2004). In motile cells, various types of actin structures were described at the cell front (e.g., branched networks in lamellipodial protrusions), and rear (mostly occupied by contractile networks), with gradients of components such as molecular motors generating flows at the scale of the entire cell (Bergert et al., 2015; Callan-Jones and Voituriez, 2016; Liu et al., 2015; Ruprecht et al., 2015; Yolland et al., 2019). Actomyosin structures described in cells and *in vitro* display striking similarities in composition and dynamic behaviour, leading to the general belief that cellular locomotion can be understood from current *in vitro* studies. Indirect methods such as speckle microscopy provided key insights into actin dynamics in living cells (Gardel et al., 2008; Hu et al., 2007; Ponti et al., 2004; Waterman-Storer et al., 1998), focusing on lamellipodial protrusions. These protrusions have been described in great detail because they are very thin, can be imaged at high resolution, and acquire simple shapes. Moreover, the preponderance of rigid 2D substrates in migration studies has historically favoured the study of adhesive lamellipodial migration over other modes such as amoeboid migration of immune and cancer cells through dense matrices.

Two major aspects have limited the study of dynamic actomyosin structures in the cortex of living amoeboid cells: 1) the network density is usually too high to observe individual filaments; 2) the large diversity of actin structures associated with various parts of the cells is hard to relate to the complex shapes adopted by motile cells to provide an integrated description of locomotion at the cell level. To understand the complex molecular machinery of cell motility in space and time, it is necessary to develop minimalistic models of amoeboid cell migration, amenable to high spatiotemporal imaging and sophisticated statistical analysis of actomyosin dynamics. Simpler actin networks are observed in amoeboid cells migrating in the absence of focal adhesions and stress fibres (Friedl et al., 1998; Lämmermann et al., 2008; Liu et al., 2015; Paluch et al., 2016; Ruprecht et al., 2015). This type of migration requires specific environments providing confinement to the motile objects, which allows propulsion by either friction or normal forces (Bergert et al., 2015; Le Berre et al., 2013; Malawista and Chevance, 1997; Reversat et al., 2020). Motile cell fragments, which are devoid of most large organelles like the nucleus but display robust migration, have been used to study lamellipodial based migration (Euteneuer and Schliwa, 1984; Malawista and Van Blaricom, 1987; Verkhovsky et al., 1999). Motile cell fragments can also be produced in vivo by cancer cells where they modulate the response of the immune system to the presence of the tumour (Headley et al., 2016). In these cells, fragments form from blebs and display rounded shapes reminiscent of amoeboid cells. Confined non-adhesive migration of bleb-based motile cell fragments thus appears as a physiologically relevant minimal motile system to study amoeboid migration.

Here, we combine confinement of living cells to induce the formation of stable blebs - motile structures with a simple shape - with high numerical aperture total internal reflection fluorescence (hNA TIRF) microscopy. hNA TIRF allowed us to investigate the assembly of the flowing actin cortex from single actin filaments at 1s time resolution over minutes, to the whole cell. It revealed that the actin cortex behaves as a passive solid advected by the rear-located myosin motors. This rigid state appears with a sharp transition at the leading edge in a low actin density bleb like protrusion. We propose a physical model that shows how a selforganized spatial patterning of the actin filaments network emerges due to the active advection by motors, combined with a rigidity percolation transition. This spatial organization determines the stability and mechanical properties (soft front and solid back) of the network. Embedding motile blebs inside collagen matrices, we show that advected percolation constitutes a robust amoeboid locomotion mechanism. Our findings explain, from first principles, the seemingly complex and fascinating capacity of an amoeboid cell to rapidly change shape and efficiently navigate complex environments.

## Results

### Characterisation of bleb fragments as a minimal amoeboid motile system

To study the dynamics of actomyosin in the cortex in the context of amoeboid migration, we chose to use 2D confined motile blebs (Liu et al., 2015; Logue et al., 2015; Ruprecht et al., 2015). We used HeLa-Kyoto cells expressing Lifeact-mCherry and MYH9-GFP (non-muscle myosin IIA, NMIIA), confined them to 3 μm and first imaged them with spinning-disc confocal microscopy. Confinement induced the formation of large and stable blebs, as described before for many cell types from different organisms (Brunet et al., 2021; Liu et al., 2015; Logue et al., 2015; Ruprecht et al., 2015). Upon confinement, canonical transient round blebs initially formed (Charras et al., 2008), followed, after about 15 s, by the appearance of blebs with an elongated shape and a longer lifetime (Fig. 1A-B and Movie S1). The time of formation of elongated blebs corresponded to the recruitment of NMIIA to the cell cortex upon confinement and the increase in the outward pushed force produced by the cell due to increased contractility (Fig. S1A-E) (Lomakin et al., 2020). As shown previously, it depended on cPLA2 and intracellular calcium, assessed by pharmacological inhibition with AACOCF3 and 2-APB respectively (Fig. S1F-J) (Lomakin et al., 2020). Some of the stable blebs eventually formed larger protrusions displaying characteristic features of motile leader blebs (Liu et al., 2015; Logue et al., 2015; Ruprecht et al., 2015), i.e. a stable actomyosin gradient and a retrograde flow (Fig. 1C). When cells were plated on poly-L-lysine, confinement induced stable blebs similarly to PEG coating, but a fraction of them moved away from the immobile cell body and eventually detached (Fig. 1D-E and Movie S1). These cytokineplasts, like keratocyte-derived fragments (Verkhovsky et al., 1999), displayed persistent motility and maintained a simple neutrophil-like amoeboid shape as they moved (Fig. 1F and Movie S1). They also maintained the same actomyosin organization observed in stable blebs - their leading edge appeared mostly devoid of actin filaments. Overall, these observations show that cell confinement and induced cortical actomyosin contractility can lead to the formation of persistent amoeboid cytoplasmic fragments with simple shapes displaying a low actin density at their front and autonomous motility. This suggests that they constitute an appropriate minimal amoeboid motile system to study the dynamics of the actin cortex.

**Figure 1:**
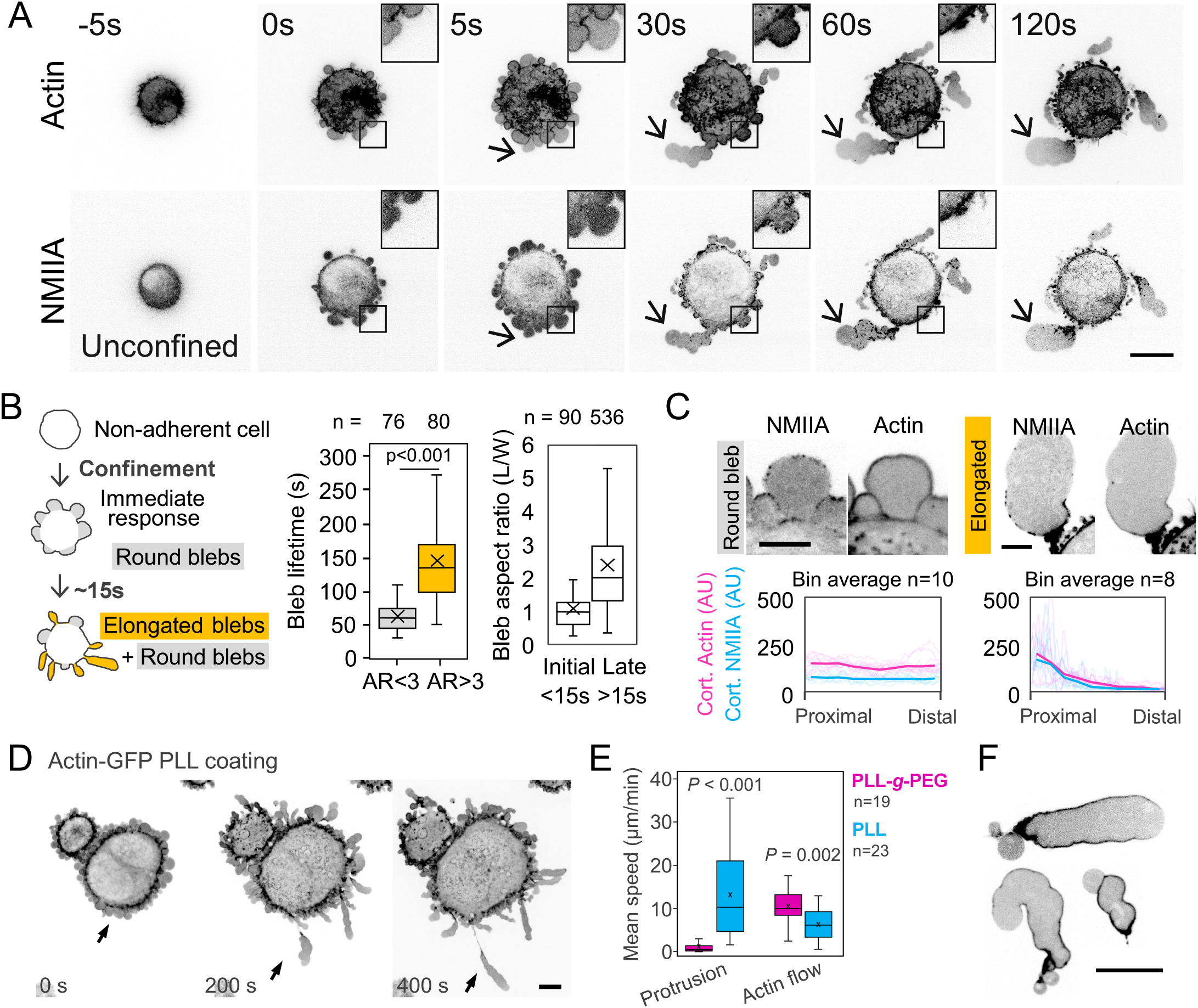
Cell confinement leads to the formation of motile fragments. **A:** Representative confocal fluorescence microscopy time-lapse images of HeLa Lifeact-mCherry (Actin, top row) *MYH9-eGFP* (NMIIA, bottom row) confined to 3μm inside a nonadhesive 2D confiner device. Time stamp, time from confinement in seconds. Square, magnification of a transient bleb protrusion-retraction cycle. Arrow, formation of a stable bleb. Scale bar, 10 μm. **B:** Left, schematic representation of the morphological features of blebs formed upon confinement. The immediate response of non-adherent cells to confinement is the formation of round, transient blebs, represented in grey. After 15 seconds approximately, elongated stable blebs, represented in orange, start to appear. Centre, individual bleb lifetime (the difference between the time of formation and time when the bleb is completely retracted, Welch’s *P*<0.001) for round (n = 76) and elongated blebs (n = 80). Right, quartile intervals and mean (cross) of the aspect ratio (length/width) for all blebs in the first few time points (15s) versus all blebs after the first time points. For clarity, the boxplot excludes the outliers. **C:** Top, representative fluorescence confocal images of a round (Round bleb, grey, left column) and an elongated bleb (Elongated, orange, right column) from 3 μm-confined HeLa *MYH9*-eGFP (NMIIA, left image) Lifeact-mCherry (Actin, right image). Scale bar, 5 μm. Bottom, averaged cortical actin (magenta) or NMIIA density (cyan), binned and normalized by position along bleb length, for representative round (n = 10) and elongated (n = 8) blebs. Shaded curves, individual data. **D:** Representative fluorescence confocal time-lapse images of HeLa *ActB*-GFP cells confined to 3μm inside a PLL-coated 2D confiner device. Time stamp, time from confinement in seconds. Arrow, stable bleb separating from the main cell body. Scale bar, 10 μm. **E:** Boxplot of front protrusion and actin flow average speed in stable blebs inside a PLL-*g*-PEG-coated microfluidic chamber (magenta, “PLL-*g*-PEG”), or inside a PLL-coated microfluidic chamber (cyan, “PLL”) (box line, median; box cross, mean; n PLL-*g*-PEG = 19 cells, n PLL = 23 cells; *P* value, Welch’s *t* test). **F:** Representative fluorescence confocal images of cytoplasts originated from HeLa *ActB-GFP* blebs confined to 3μm inside a PLL-coated (adhesive) microfluidic chamber. Scale bar, 10 μm.

### Imaging single actin filaments in live stable bleb protrusions

Because spinning disc imaging was not sufficient to resolve single actin filaments even in the low filament density region at the tip of the stable blebs, we turned to high numerical aperture total internal reflection fluorescence (hNA TIRF) microscopy. Since large stable blebs develop between two confinement slides, their membrane is pressed against the bottom coverslip, making TIRF imaging possible over the broad cell-substrate contact area from the cell front to the rear. We first compared HeLa cells expressing either lifeact-GFP, f-tractin-GFP and a HeLa cell line with TALENS-edited actin-GFP in the actin locus. As a control, we also fixed cells under confinement and stained them with phalloidin. While the general picture was similar for all three cell lines, the actin-GFP cells displayed a much better contrast, allowing the visualization of the network at the actin filament level at the bleb front (Fig. S2A-F). Protruding regions appeared first totally devoid of actin filaments as expected for an early blebbing membrane and were populated within a minute by short filaments of 0.65 ± 0.45 μm length (Fig. 2A-B and Movie S2). Protruding regions maintained a sparse density network (1.19 ± 0.29 μm actin filaments per μm^2^) with low branching density (0.49 ± 0.14 branches per μm of filament). Unconnected short filaments displayed unbiased, seemingly diffusive, mostly rotational motion, and stayed on the focus of the TIRF plane, suggesting they were loosely attached to the bare membrane. The diffusive motion stopped in filaments forming clusters (Fig. 2C-D). Single actin filament branching had never been directly imaged in live cells. We thus characterized it by measuring the elongation speed (0.25 ± 0.24 μm s^-1^, Fig. 2E and Movie S2), in the order of 1 μm s^-1^ as previously inferred from indirect methods (Funk et al., 2019; Pollard and Mooseker, 1981) and the branching angles (65 ± 10°, Fig. 2F and Movie S2), which appeared close to 70° as expected (Svitkina et al., 1997). Seeds and branches often collapsed onto pre-existing actin filaments to form bundles (Fig. 2G-H and Movie S2). By quantifying these features, we concluded that we were able to resolve single actin filaments and that protruding parts of stable blebs display a low density of short freely diffusing and elongating actin filaments. This region of freely diffusing motion of low-density actin filaments can be described as a gas-like state of the actin network.

**Figure 2:**
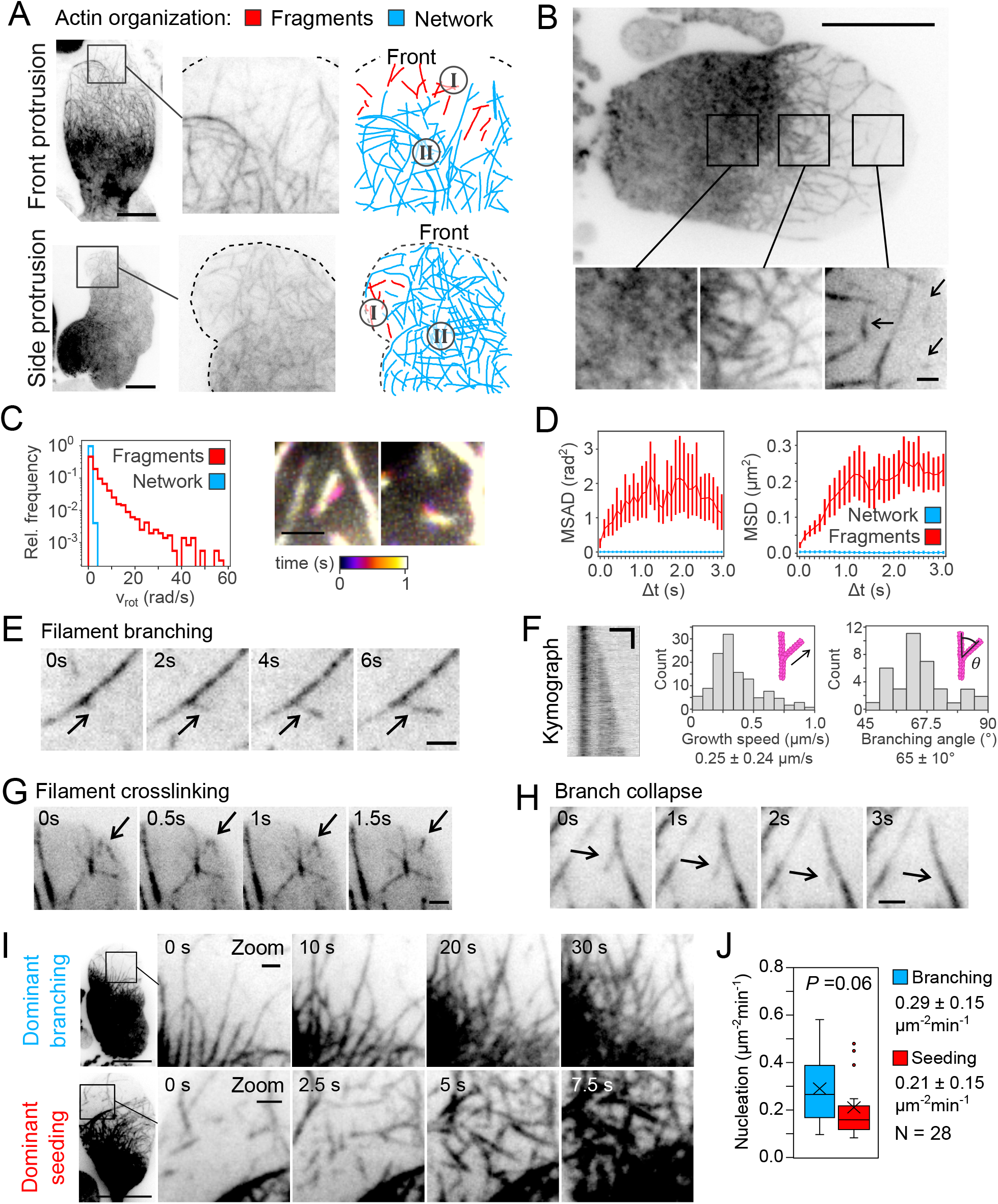
Actin filament dynamics in stable blebs. **A:** Left and middle panels, representative high numerical aperture TIRF images of 3 μm-confined HeLa *ActB*-GFP cells. Right panels, analysis of the actin network. Top panels, stable bleb protruding at the front. Bottom panels, stable bleb protruding to the left side. Red lines and label “I”, actin filaments not linked to the main actin network. Blue lines and label “II”, main actin network. Scale bar, 10μm. **B:** Top, representative high numerical aperture TIRF image of a stable bleb from 3 μm-confined HeLa *ActB*-GFP cells. Scale bar, 10 μm. Bottom, zoom of the regions marked with black squares. Arrows, actin filaments not connected to the main network. Scale bar, 1 μm. **C:** Left, histogram of the rotational velocity (*v_rot_*, rad/s) of actin filaments not connected to the main network (fragments, red), or connected to the network (network, cyan). Right, representative high numerical aperture TIRF color-coded time projections of actin filaments not connected to the main network, in 3 μm-confined HeLa *ActB*-GFP cells. Scale bar, 1μm. **D:** Left, mean square angular displacement (MSAD, rad^2^) of actin filaments not connected to the main network (fragments, red), or connected to the network (network, cyan). Right, mean square displacement (MSD, μm^2^) of actin filaments not connected to the main network (fragments, red), or connected to the network (network, cyan). **E:** Representative high numerical aperture TIRF time-lapse images of a filament elongation event, marked with black arrows. Scale bar, 1μm. **F:** Left, kymograph of the branching event from panel 2E. Scale bar, 1μm/s. Right, histogram of filament branching elongation speed (μm/s) and branching angle (°). Label, mean value ± standard deviation. **G:** Representative high numerical aperture TIRF time-lapse images of a filament crosslinking event, marked with black arrows. Scale bar: 1μm. **H:** Representative high numerical aperture TIRF time-lapse images of a branch growth and collapse event, marked with black arrows. Scale bar: 1μm. **I:** Representative high numerical aperture TIRF time-lapse images of a bleb with network branching (top) or network seeding (bottom) as main assembly mechanism. **J:** Frequency of nucleation of network segments by branching (cyan) or seeding (red), measured as number of events per μm^2^min at the bleb front.

### Assembly of the actin cortex from single filaments at the tip of stable blebs

To examine how the region with a low density of filaments at the bleb tip became populated by a dense actin network, we performed imaging at high temporal resolution. The appearance of new actin seeds and the branching from existing filaments contributed similarly to the formation of new network segments (probability of branch nucleation: 0.29 ± 0.15 μm^-2^min^-1^, probability of seed nucleation: 0.21 ± 0.15 μm^-2^min^-1^, Fig. 2I-J). This detailed observation of the bleb front suggests that bundling of actin filaments is an important driver of the actin network assembly, consistent with the known requirement for actin crosslinkers for stable bleb formation (Logue et al., 2015). We thus imaged cells expressing the actin filaments bundler alpha-actinin (Charras et al., 2006) fused to mCherry in addition to actin-GFP and performed two-colour hNA TIRF imaging. This enabled to clearly distinguish single actin filaments, devoid of alpha-actinin-mCherry staining, and regions of filaments overlap, which displayed both higher actin-GFP staining and were positive for alpha-actinin-mCherry (Fig. S2G-H and Movie S2). Overall, these observations suggest that the protruding region at the bleb tip is almost devoid of actin filaments and gets populated by a crosslinked actin network mostly assembled from short elongating and branching filaments that diffuse and bind to each other. The low density of actin filaments explains the lack of force transmission through the network to the front bleb membrane and thus explains the lack of retraction and the stability of the bleb front.

### Retrograde advection of a non-contractile crosslinked actin network maintains a low actin density front and ensure bleb stability

We then asked how a low density of actin filaments is maintained at the bleb front despite the constant appearance and elongation of new filaments. The stability and the locomotion capacity of leader blebs were attributed to the generation of a steady retrograde actin flow produced by NMIIA motors (Callan-Jones and Voituriez, 2016; Liu et al., 2015; Ruprecht et al., 2015), visible in both the spinning disc confocal and in the hNA TIRF. This flow is the hallmark of prototypic amoeboid cells. It was proposed to have 3 essential functions: i) depletion of actomyosin front the front leading to a pressure-driven balloon-shaped front, ii) creating a steady-state contractility gradient with higher contractility at the rear, and iii) generating the propulsion force via coupling non-specific frictional molecules with the substrate (Callan-Jones and Voituriez, 2016; Liu et al., 2015; Ruprecht et al., 2015). To investigate the formation of the flowing actomyosin network at high spatiotemporal resolution we imaged actin-GFP and NMIIA-mApple expressing cells with hNA TIRF microscopy. Unexpectedly, we found that NMIIA was strictly located at the very rear of the bleb, while actin was reaching closer to the front tip (Fig. 3A and Movie S3, single example, and Fig. 3B quantified on several blebs and time points). Closer examination of the NMIIA images showed that even very small NMIIA clusters did not reach towards the front of the bleb and appeared only in the already dense actin region, then forming larger clusters towards the bleb rear (Fig. S3A). This suggested that a large fraction of the dense actin cortex region was formed only of passive crosslinked actin filaments without any contractile activity. To confirm that, we analysed the dynamics of actin and myosin using particle image velocimetry (PIV). Visual examination of the actin velocity profile showed an isotropic, disordered slow velocity field at the front, consistent with unbiased diffusing filaments, followed by a fast quasi-uniform coherent retrograde flow in the middle and a slower convergent flow (divergence 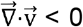) at the rear of the bleb (Fig. 3C). This was confirmed by an analysis of the divergence and average velocity (Fig. 3C, single example, and Fig. 3D quantified on several blebs and time points). Combined with the density map, the flow map allowed to establish actin and NMIIA net turnover maps (Fig. 3E, single example, and Fig. 3F quantified on several blebs and time points), which confirmed the visual examination: while actin filaments assembled on almost the whole length of the bleb and disassembled at the very rear, NMIIA recruitment was shifted to the rear. Further analysis of the actin flow field showed a large region in the middle of the bleb with aligned velocity vectors parallel to the bleb long axis, displaying a large correlation length (Fig. 3G, single example, and Fig. 3H quantified on several blebs and time points). Overall, this analysis suggests the existence of a non-contractile crosslinked actin network in the middle of the bleb, which was not anticipated from previous studies of actin flows in stable/leader blebs. This passively advected strain-free actin network bridges the dispersed gas-like cortex at the bleb front to an active contractile network at the bleb rear (Fig. S3B).

**Figure 3:**
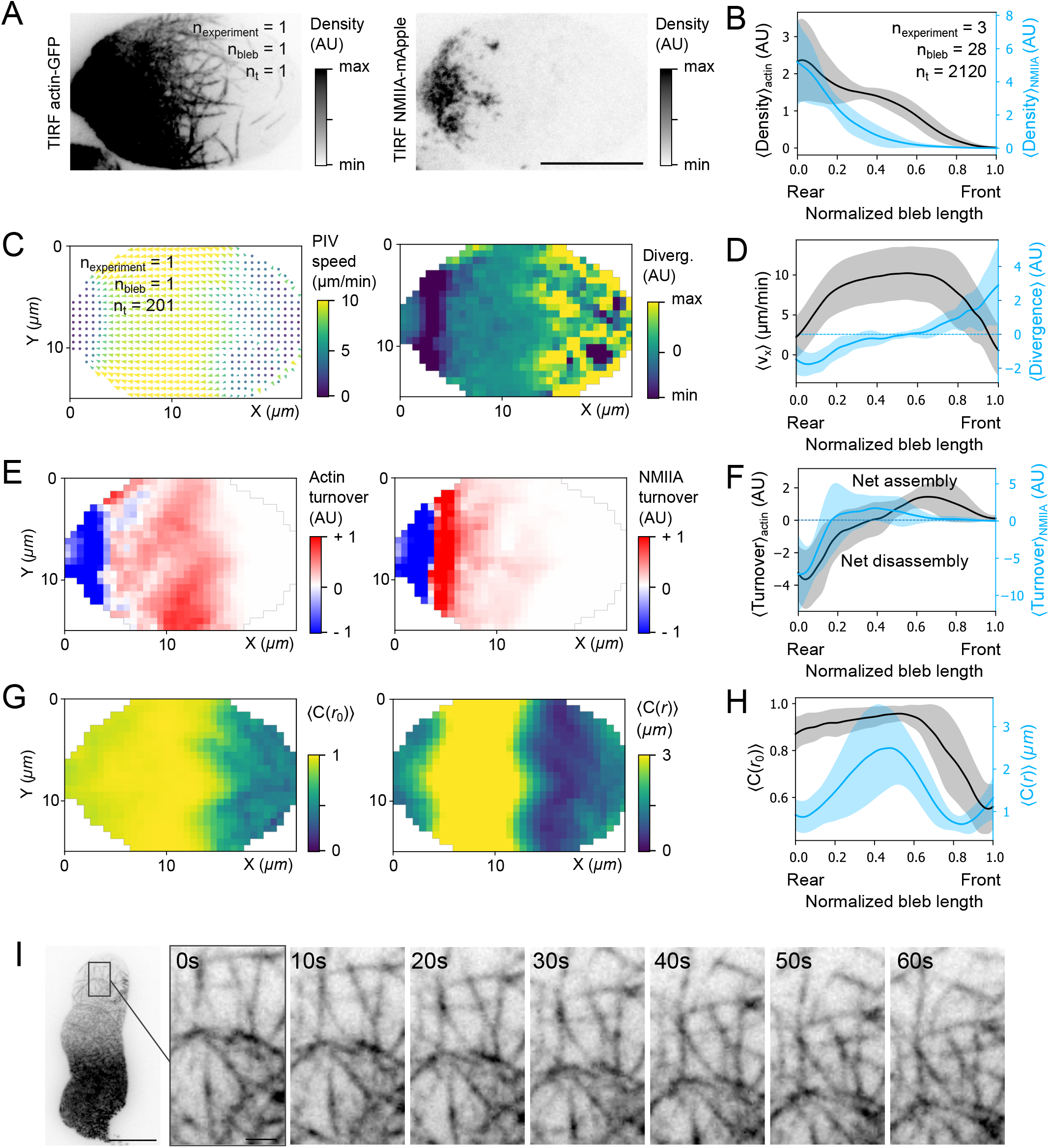
Actomyosin network assembly and dynamics in stable blebs. **A:** Representative hNA TIRF image of 3 μm-confined HeLa *ActB*-GFP (TIRF actin-GFP, left) cells transfected with mApple-MyosinIIA (TIRF NMIIA-mApple, right). Scale bar, 10 μm. **B:** Actin (black, mean ± s.d.) and NMIIA (blue, mean ± s.d.) density profile along bleb length, averaged over the width, time, and several blebs (N = 3, n_bleb_ = 20, n_t_ = 2120). **C:** Left, time-averaged PIV of a single bleb (n_bleb_ = 1, n_t_ = 201). Right, time-averaged PIV divergence field of a single bleb (nbleb = 1, n_t_ = 201). **D:** PIV horizontal component (v_x_, black, mean ± s.d.) and PIV divergence (blue, mean ± s.d.) along bleb length, averaged over the width, time, and several blebs (N = 3, n_bleb_ = 20, n_t_ = 2120). **E:** Left, time-averaged net actin turnover of a single bleb (n_bleb_ = 1, n_t_ = 201). Right, time averaged net NMIIA turnover of a single bleb (n_bleb_ = 1, n_t_ = 201). **F:** Net actin turnover (black, mean ± s.d.) and net NMIIA turnover (blue, mean ± s.d.) along bleb length, averaged over the width, time, and several blebs (N = 3, n_bleb_ = 20, n_t_ = 2120). **G:** Left, time-averaged local correlation C(r_0_) of a single bleb (n_bleb_ = 1, n_t_ = 201). Right, time-averaged correlation length *C*(*r*) of a single bleb (n_bleb_ = 1, n_t_ = 201). **H:** Local alignment *C*(*r*_0_) (black, mean ± s.d.) and correlation length *C*(*r*) (blue, mean ± s.d.) along bleb length, averaged over the width, time, and several blebs (N = 3, n_bleb_ = 20, n_t_ = 2120). **I:** Left, Representative high numerical aperture TIRF time-lapse images of 3 μm-confined HeLa *ActB*-GFP cells. Scale bar, 10 μm. Right, time-lapse images of a bleb front region.

We then asked what is the state of the passive actin network located in the elongated flat region of the bleb, between the low actin density front and the contractile back of the bleb. The PIV analysis showed a long-range correlation in the flow in the central region of the bleb, demonstrating the absence of contractile activity, but the actin filaments in this region could be flowing either in a fluid- or in a solid-like state. By definition, in a solid, conversely to a fluid, the relative position of constituents does not change over time. We first followed the relative position of specific actin structures at the single filament scale within the network. We found that, in the region devoid of NMIIA, landmark structures in the actin cortex were advected backwards without showing any deformation (Fig. 3I, red square, and Fig. S3C, Movie S3), and noticed that this solid-like state of the actin network corresponded to the flatter part of the elongated blebs (i.e. the contour curvature approaching zero, Fig. S3D black line, contour curvature). This suggests that the overall elongated shape of stable blebs might be due to the presence of an intermediate non-contractile solid actin network between the front and the rear. Since actomyosin networks are known to have a visco-elastic nature, the absence of relative motion between the elements of the network could be due to very low forces exerted on the network (for example due to the absence of adhesion to the substrate) and to the short timescale of the observation (due to the fast speed of the flow). We thus can conclude that, at the timescale and in the range of forces involved in the migration of stable blebs, the central part of the actin cortex effectively behaves as a solid-like network. This is further supported by the observation of local kinks on the bleb outline, formed due to fluctuations of the front membrane and stabilised by the actin cortex assembly. These kinks moved towards the rear keeping an identical shape (Fig. S3C, arrow and star), further confirming that this part corresponded to a rear-advected strain-free solid network. Together these observations support the existence of a passive solid-like crosslinked actin network implicated in the shape of stable motile blebs and occupying most of the bleb length.

### A rigidity percolation transition in the actin network explains the sharp transition between the gas-like and the solid-like phase at the bleb front

The transition between a diluted (gas-like) state and a solid state is reminiscent of the sol-gel transition model initially proposed to explain the motility of amoeboid cells (Hyman, 1917; Mast, 1932; Pantin, 1923). However, while gelation describes the transition from a solution to a gel, the gel is not necessarily in a rigid solid state (e.g., if it is not crosslinked). On the contrary, our observation points to the assembly of a rigid network. Such transition is generally described as a rigidity percolation transition critically controlled by the concentration of the constituents (Alvarado et al., 2017). A network is percolated when its constituents (filaments and crosslinker) are physically connected. In the case of rigid filaments and crosslinkers, this purely geometric definition coincides with the mechanical definition of rigidity, which implies long-range stress propagation: forces applied on any part of the network (e.g., contractile forces at the rear) are transmitted to all the parts (e.g., the front edge of the network). In the case of stable blebs studied here, contrary to a classical bulk percolation transition with a uniform actin concentration, we propose that the rigidity percolation transition is spatially organized because motors generate a gradient of actin filament concentration by advecting the percolated network.

To test whether the transition between the gas-like state of the actin network at the bleb front and the advected solid-like part can be described as a percolation transition organized in space by a gradient of filament concentration, we combined several types of analysis of the actin images. First, we segmented the TIRF actin channel using a machine-learning pipeline (Berg et al., 2019) and used the skeletonized images (Fig. 4A) to identify clusters of filaments and measure the actin occupancy *p* (area fraction occupied by actin filaments). Importantly, occupancy and density were showing a similar trend until full occupancy. Density then continued to increase until the rear of the bleb, as filaments that could not be resolved continued to accumulate in the actin cortex (Fig. 4B-C). We first investigated key predictions of classical percolation theory (Stauffer et al., 1994): the existence of a threshold occupancy to reach percolation, and the relation between cluster sizes and occupancy. We identified the critical occupancy 〈*p_c_*〉 corresponding to the percolation transition (Fig. 4D-E). We also observed, at the critical occupancy, the expected behaviour of the mean size of the two largest clusters (Fig. 4F-H), and a cluster area distribution consistent with a power-law around the percolation threshold *p_c_* (Fig. 4I). Importantly, although the occupancy measured here is not the occupancy of the filaments themselves but their segmented binary image, it does not affect the characterisation of the transition, but only the numerical value of the critical density at the percolation threshold, which is expected to be system dependent. We estimated here the actin filament density at the percolation threshold to be of 2.15 ± 0.24 μm of filament per μm^-2^ of membrane. Second, we focused on the coupling between percolation and advection of the percolated clusters, a phenomenon not reported before for percolated networks. We investigated the correlation between the spatial flow correlation obtained from the PIV analysis (Fig. 4J) and the occupancy (Fig. 4K) or the actin density (Fig. 4L). The correlation length *C*(*r*) transitioned from low to high around the percolation threshold *p_c_*, while the local alignment *C*(*r*_0_) already reached the maximum at the percolation threshold *p_c_*. This suggests that at the percolation threshold *p_c_* the network shows signs of criticality – local clustering but not long-range correlation. Overall, these analyses confirm that the actin network undergoes a percolation transition organized in space by a gradient of filament concentration, leading to the formation of an actively advected rigid phase that bridges the low-density front and the contractile rear of the bleb.

**Figure 4:**
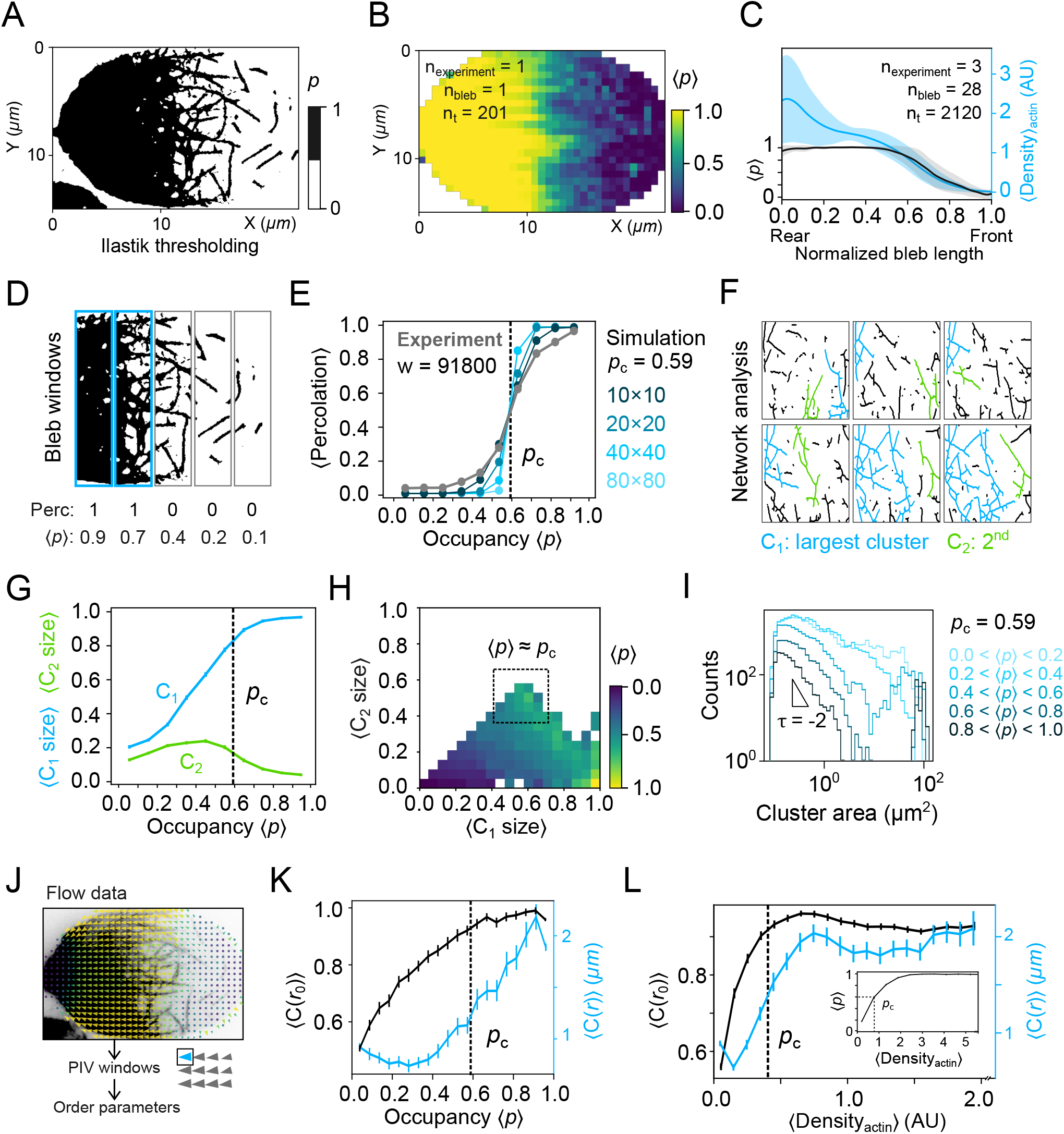
Evidence of a sol-gel percolation transition in the actomyosin network. **A:** Actin segmentation of panel 3A performed by an user-trained machine-learning algorithm (Ilastik thresholding). Black, pixels corresponding to actin. White, background. **B:** Time-averaged average occupancy 〈p〉 of a single bleb (bleb shown in panel 4A, n_bleb_ = 1, n_t_ = 201). **C:** Average occupancy 〈p〉 (black, mean ± s.d.) and actin density (blue, mean ± s.d.) along bleb length, averaged over the width, time, and several blebs (N = 3, n_bleb_ = 20, n_t_ = 2120). **D:** Schematic representation of the percolation analysis, using the bleb shown in panel 4A. Top, actin-segmented images are divided in transversal windows. Bottom, average occupancy 〈p〉 and percolation (0 for non-percolated, 1 for percolated) are measured for each window. **E:** Average percolation probability as a function of the binned average occupancy, n = 91800 bleb windows. Gray, experimental data. Black and blue shades, simulations. Simulations, average percolation probability as a function of the binned average occupancy in site-percolation simulations for different system sizes (10×10, 20×20, 40×40, 80×80). Dashed line, theoretical percolation threshold (*p_c_* = 0.59) (Stauffer et al., 1994). **F:** Schematic representation of the network analysis in panels 4G-I. Blebs are divided in windows as in panel 4D, where cluster sizes (cluster length divided by window length), cluster areas (cluster convex areas) and the window average occupancy 〈p〉 are measured. **G:** Average biggest (cyan line, C_1_) and second biggest (green line, C_2_) cluster sizes, normalized to system length, as a function of the binned average occupancy, n = 91800 bleb windows. Dashed line, theoretical percolation threshold (p_c_ = 0.59). Bars, 95% confidence interval. **H:** Binned average occupancy as a function of the biggest and second biggest cluster sizes relative to system length, n = 91800 bleb windows. Dashed square, region where both the biggest and the second biggest cluster sizes approach the system length. **I:** Log-scaled histogram of cluster areas in windows grouped by average occupancy. Colours, average occupancy. Label, theoretical square site percolation threshold (p_c_ = 0.59). **J:** Schematic representation of the analysis of panels 4K-L. Order parameters are plotted as a function of average occupancy 〈p〉 and actin density, calculated for each PIV window. **K:** Average local correlation *C*(*r*_0_) (black) and correlation length *C*(*r*) (blue) as a function of the binned average occupancy. Dashed line, theoretical percolation threshold (p_c_ = 0.59). Bars, 95% confidence interval. **L:** Average the local alignment *C*(*r*_0_) (black) and correlation length *C*(*r*) (blue) as a function of the binned average actin density. Dashed line, density corresponding to the theoretical square site percolation threshold (Density_actin_ = 0.4). Bars, 95% confidence interval. Inset, average occupancy as a function of the binned average actin density. Dashed line, theoretical percolation threshold p_c_ = 0.59), and corresponding density value (Density_actin_ = 0.4). Bars, 95% confidence interval.

### The percolated actin network ensures a direct force transmission from the contractile rear to the front membrane of the bleb

We then asked how the gradient of actin filament concentration drives force transmission. In previous works, it has been proposed that actomyosin assemblies can self-organize and produce stable flows and concentration gradients (Ierushalmi et al., 2020; Ruprecht et al., 2015). In these studies, actomyosin was treated as a continuous viscoelastic fluid, while our high NA TIRF imaging revealed that the actin filaments are advected as a rigid network. In contrast to classical active fluid models, where actin flows and stress decay gradually away from contractile regions, the consequences of the rigidity percolation transition are the long range propagation of stress in the percolated cluster, and the sharp transition from uniform flow to no flow below the percolation threshold *p_c_*. We were able to indirectly visualize the forces transmitted through the crosslinked network via their effect on the re-orientation of pulled filaments and network rupture events (Fig. 5A-B and Movie S4). Forces applied on the percolated network were also transmitted to the bleb membrane, when percolated actin filaments bound to the flowing network also connected the free membrane at the front. These events produced kinks in the membrane and were followed either by i) a detachment of the filament, leading to snapping of the membrane away from the actin network and recovery of the membrane shape (Fig. 5C and Movie S4) or ii) retraction (Fig. 5D and Movie S4), reminiscent of what has been observed in reconstituted systems (Litschel et al., 2021). Together these results show that the percolated actin network, although it does not contain molecular motors, transmits forces on long ranges at the scale of the entire bleb (which can reach several tens of microns in length), ensuring a mechanical coupling between the force-producing actomyosin network at the rear and the front of the bleb.

**Figure 5:**
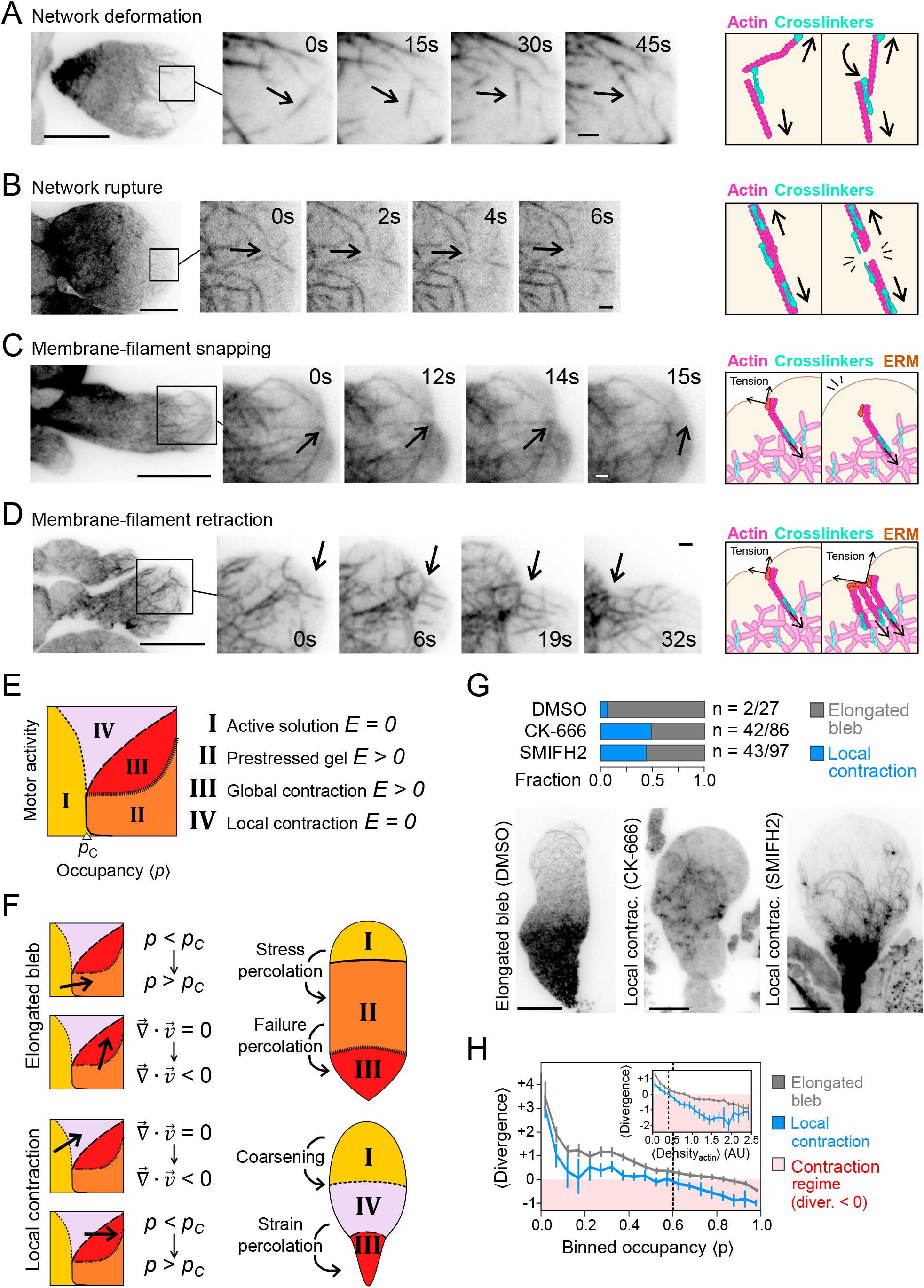
Force transmission through the actomyosin network. **A:** Left, representative high numerical aperture TIRF time-lapse images of HeLa *ActB*-GFP cells displaying network deformation (black arrows). Scale bar: 10μm (left, whole bleb) and 1μm (right, time-lapse sequence enlarged image from black square). Right, schematic representing the event. **B:** Left, representative high numerical aperture TIRF time-lapse images of HeLa *ActB*-GFP cells displaying network rupture (black arrows). Scale bar: 10μm (left, whole bleb) and 1μm (right, time-lapse sequence enlarged image from black square). Right, schematic representing the event. **C:** Left, representative high numerical aperture TIRF time-lapse images of HeLa *ActB*-GFP cells displaying membrane-filament snapping (black arrows). Scale bar: 10μm (left, whole bleb) and 1μm (right, time-lapse sequence enlarged image from black square). Right, schematic representing the event. **D:** Left, representative high numerical aperture TIRF time-lapse images of HeLa *ActB*-GFP cells displaying membrane-filament retraction (black arrows). Scale bar: 10μm (left, whole bleb) and 1μm (right, time-lapse sequence enlarged image from black square). Right, schematic representing the event. **E:** Left, non-equilibrium regimes in active systems: active solution (I), pre-stressed gels (II), global contraction (III), and local contraction (IV). Right, Young’s modulus (*E*) is greater than zero in states II and III, where long-range force transmission can occur. Diagram adapted from Alvarado et al., 2017. **F:** Top, diagram showing the spatial organization of the three cortical regimes and its correspondence to bleb shape in elongated blebs. Stress percolation: the active solution regime at the bleb front undergoes a stress percolation transition at 〈p〉 >*p_c_* to a prestressed gel regime. Failure percolation: the prestressed gel regime undergoes a failure percolation transition at the rear to a global contraction regime, at which 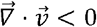. Bottom, diagram showing the spatial organization of the three cortical regimes and its correspondence to bleb shape in blebs displaying a local contraction regime. Coarsening, the active solution regime at the bleb front undergoes a coarsening transition to a local contraction regime, contracting 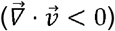 at 〈p〉 < *p_c_*. Strain percolation, the local contraction regime undergoes a strain percolation transition at the rear to a global contraction regime, at 〈p〉 > *p_c_*. **G:** Top, fraction of elongated (grey) versus blebs displaying a local contraction regime (cyan) in control (DMSO), 100 μM CK-666 and 40 μM SMIFH2 conditions. Bottom, representative high numerical aperture TIRF images of blebs from HeLa *ActB*-GFP cells displaying an elongated regime (DMSO, left), or local contraction regime (Local contrac.) under CK-666 (CK-666, middle) and SMIFH2 (SMIFH2, right) conditions. Scale bar, 10 μm. **H:** Average flow divergence as a function of the average binned actin density. Grey, average for elongated stable blebs (N = 3, n_bleb_ = 20, n_t_ = 2120). Cyan, representative example of a bleb displaying a local contraction regime, corresponding to the SMIFH2-treated bleb displayed in panel 5G. Dashed line, density corresponding to the percolation threshold (*p_c_* = 0.59). Bars, 95% confidence interval. Red region, contraction regime (divergence < 0). Inset, same data plotted as a function of the actin density instead of the binned occupancy. Dashed line, density corresponding to the percolation threshold (*p_c_* = 0.59, Density_actin_ = 0.4).

### A working model of advected percolation explains bleb stability and morphogenesis

A previous study based on *in vitro* reconstituted actin networks also proposed to use the framework of percolation theory to describe transitions in the bulk properties of the network (Alvarado et al., 2017, 2013). In this study, a phase diagram was established, with two main control parameters, motor activity and network connectivity, defining various states of the actin network (Fig. 5E). These include the states we have identified in blebs, the active solution (or gas-like state in Fig. S3B) corresponding to the bleb front, the prestressed gel (or solid-like state) corresponding to the bleb middle, and the global contraction (or contractile state) corresponding to the bleb rear (Fig. 5F top part). In contrast to bulk networks studied *in vitro*, the actin network in blebs is contained in a membrane which imposes geometric boundaries. We propose that the bleb geometry leads to the spatial self-organization of the actin network, which undergoes two consecutive transitions from the bleb front (passive gas-like phase) to the bleb middle region (advected, percolated, passive, strain-free solid network) and to the bleb rear (contractile network driving advection). The actin network in the bleb follows a well-defined path through the phase diagram, from its front to its rear, first increasing in occupancy until crossing the percolation threshold (*p_c_*) and then increasing in motor activity, generating a global contraction and flow (Fig. 5F top part). Such selforganization in space requires the spontaneous establishment of both a gradient of actin filaments and a gradient of myosin. We argue that such gradients can be self-induced by the advection of the network (due to the myosin motors accumulation at the rear), similar to the previously proposed “universal coupling between speed and persistence” model UCSP model, which shows how actomyosin flows can feedback on cell polarity (Maiuri et al., 2015). This results in the stable coexistence of the previously described bulk phases along the length of the bleb, explaining the bleb stability and shape.

To further test this working model, we perturbed the actomyosin network. Previous studies reported the drastic effect of inhibiting actin turnover (treatment with Jasplakinolide) and myosin motors (treatment with Blebbistatin) and found that these treatments rapidly lead to the loss of the stable bleb state (Liu et al., 2015; Ruprecht et al., 2015). We thus applied a milder perturbation and reduced the rate of actin nucleation using drugs against the main actin nucleation factors, formins and the Arp2/3 complex. Because the percolation threshold is reached for a critical actin density allowing most filaments to establish physical connections, reducing the rate of actin nucleation should perturb the position of the percolation transition and thus the bleb shape. Treating cells with either formin or Arp2/3 inhibitors had a similar effect. 60 minutes after the start of the treatment, a fraction of blebs appeared more spherical and the transition from the low density to the high-density actin network occurred further towards the rear of the blebs (Fig. 5G). Analysis of the actin dynamics showed that the onset of contraction occurred on a non-percolated network (Fig. 5H), generating characteristic aster-like patches of actin filaments, already reported before on the cortex of cells treated with actin-depolymerizing drugs (Luo et al., 2013). This suggests that the percolated non-contractile network was absent from these blebs, correlating with the loss of the elongated shape (Fig. 5G). The round shape of blebs in treated cells suggests that no force was transmitted from the contractile dense rear network to the bleb front. Indeed, the network appeared mostly fragmented. After few hours of confinement under drug treatment, the blebs eventually collapsed, and no new stable blebs were formed (Movie S5). These results suggest that a large enough rate of actin filament assembly is required to ensure that the percolation transition occurs before the onset of contractility, leading to an elongated stable bleb shape. They also support our working model of self-organisation of an advected percolated actin network.

### A physical model of advected percolation predicts the stable self-organisation of the actin network into regions of different rheological properties along the bleb length

Our working model postulates that the shape and stability of motile blebs emerge from the combination of two simple phenomena: (i) the spontaneous establishment of a gradient of actin filaments and myosin motors and (ii) a rigidity percolation transition at a critical density of actin filaments. We postulate that the combination of these two simple phenomena would lead to the spontaneous and stable appearance of spatially organized physical properties in the actomyosin network (Fig. 5F). To provide a rigorous demonstration that these ingredients are sufficient to explain our observations, we propose the advected percolation model, based on simple self-organization mechanisms. The essential ingredients of the model are: (i) a rigidity percolation transition in the actin network, which we show is coupled with (ii) the emergence of spontaneous flows in actomyosin systems.

In brief, the advected percolation model (see Supplementary Information for more details) considers a 2-dimensional cortex with nucleation (controlled by an effective rate *β* that depends linearly on the local concentration of nucleators, *c_N_*) and depolymerization of actin filaments with a rate *k_d_* (Fig. 6A). The bleb rear is denoted by *x* = 0 and the bleb front by *x* = *L*. Free actin filaments diffuse and irreversibly and rigidly bind together upon encounter, corresponding to the observation of a bleb front with actin crosslinkers but devoid of myosin II. The main cluster is defined by the set of all filaments connected to *x* = 0 and extends to *x* = *L_C_*, where the local filament concentration reaches the percolation threshold *p_c_*. We assume that both myosin motor concentration *c_M_* and actin nucleator concentration *c_N_* follow classical advection-diffusion equations, which lead to exponential steady-state profiles *C_M_*~*e*^−(*x/L_M_*)^ and *C_N_*~*e*^−(*x/L_N_*)^ respectively, where *L_M_* and *L_N_* are the typical decay length of the two profiles. In particular, this implies a similar profile of the actin assembly rate that we write as *β* = *β*_0_*e*^−(*x/LN*)^. Following observations (Fig. 1C and Fig. 3B), we take *L_M_* ≪ *L*, and *L_M_* thus defines the extension of the contractile region. We assume that the resulting gradient of contractility induces the reward advection of the main cluster (considered as a solid) at a constant speed *v*_0_ (whose value is set by the gradient of contractile stress and friction and is not determined here). For *x* < *L_M_*, the main cluster contracts due to active contractile stresses although this is not explicitly described in the model. For *L_M_* < *x* < *L_C_*, the cluster is a strain-free passive solid advected at speed v_0_. Below the critical concentration *p_c_* (i.e., typically for *x* > *L_C_*), the actin filaments are not transported and only diffuse. Analysis of our model by 2D numerical simulations (Fig. 6B-D and Fig. S4C for a simpler 1D version) and mean-field arguments (Fig. S4A,B) shows that *L_C_* is critically controlled by actin assembly/disassembly rates and advection speed and can be written *L_C_*~(*D_a_* / *v*_0_)In(*β*_0_ / *k_d_*), where *D_a_* is the effective diffusion coefficient of actin nucleators. The model recapitulates the observed phenomenology and spatial organization of blebs and shows that it is controlled by the relative magnitude of the lengths *L_C_, L_M_, L*. For *L_M_* < *L_C_* < *L* (large advection and/or moderate actin assembly), the model yields stable, persistent blebs (Fig. 6C-D, non-percolated). For *L_C_* > *L* (small advection and/or fast actin assembly) the solid cortex rapidly fills the entire bleb, which implicitly leads to the irreversible retraction of the bleb (Fig. 6C-D, percolated). For *L_C_* < *L* but *L_C_* close to *L*, we obtain intermittent dynamics, with localized, transient contacts of the main cluster with the bleb front edge made possible by fluctuations of *L_C_* (Fig. 6C-D, intermittent). Finally, this analysis predicts that for *L_C_* < *L_M_* (small *β*_0_ in the model, realized experimentally by perturbations of formins and Arp2/3 in Fig. 5E-H), the strain-free advected region disappears, leading to a fluid contractile region without stress propagation (for *L_C_* < *x* < *L_M_*), next to an active contractile solid region (*x* < *L_C_* < *L_M_*), as observed experimentally. The advected percolation model thus captures well the main features we observed experimentally: the transition between the two main types of blebs, transient and stable, the actin gradient in stable blebs, and the various protrusion dynamics. We conclude from the analysis of this physical model that the simple ingredients we implemented to construct it (rigidity percolation transition coupled with spontaneous actomyosin flows) are sufficient to explain the formation of an actin cortex with a stable spatial pattern of rheological properties from the front to the rear of elongated blebs.

**Figure 6:**
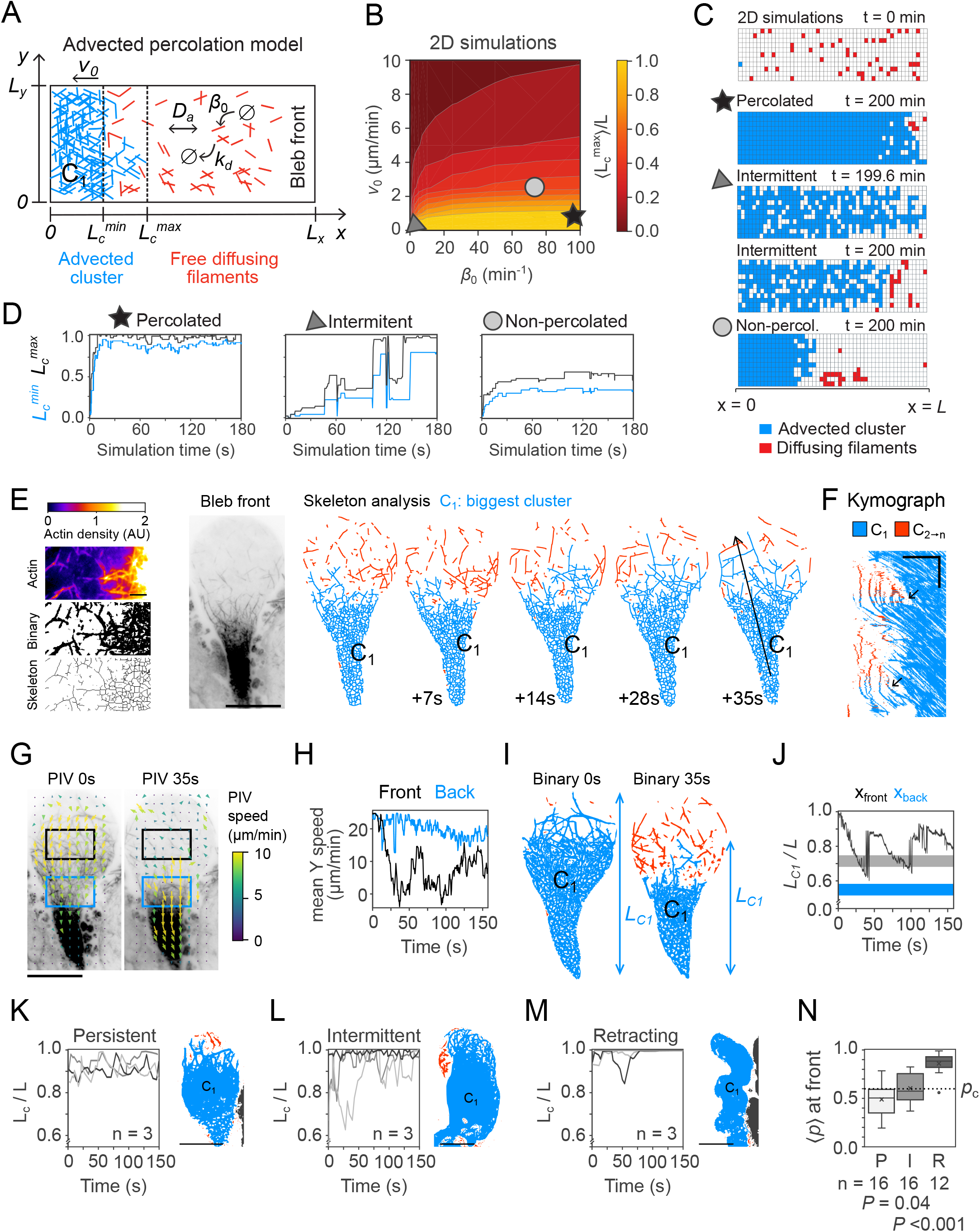
The advected percolation model. **A:** Diagram representing the advected percolation model. Parameters are defined in the text. Blue, main cluster. Red, freely-diffusing filaments, not connected to the main cluster. **B:** Outcomes of the 2D stochastic advected percolation model, phase diagram of the average maximum cluster length along the x-axis, 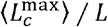, as a function of the assembly rate *β*_0_ and the advection velocity *v*_0_. Legend is shown as inset. Parameters are *L_x_* = 50 μ*m*, *L_y_* = 10μ*m*, *D_a_* = 10 μ*m*^2^*min*^-1^, *k_d_* = 0.5 *min*^-1^. **C:** Snapshots of the configurations at different time points for the three different simulations pinpointed in panel 6B (star, percolated regime; triangle, intermittent regime; circle, non-percolated regime). Blue, sites belonging to the cluster attached to the left hand-side boundary. Red, all other sites with a non-zero number of molecules. White, sites without any molecule. Time stamp, simulation time from start. **D:** Snapshots of the time trajectories of the renormalised maximum cluster length and minimum cluster length for the three different conditions pinpointed in panel 6B (star, percolated regime; triangle, intermittent regime; circle, non-percolated regime). Time, simulation time from start. **E:** Left, example network analysis of a bleb region, from the actin fluorescence channel (top) to the binarized image (middle), and the extracted skeleton (bottom). Scale bar, 1μm. Middle, representative high numerical aperture TIRF image of HeLa *ActB*-GFP cells. Scale bar, 10μm. Right, actin network analysis of skeletonized time-lapse images. Blue, labelled C_1_, largest cluster. Red, all other clusters. Time stamp, seconds elapsed from first image in the shown series. Black arrow, region plotted in the histogram in panel 6F. **F:** Kymograph of the region marked with a black arrow in panel 6E. Blue, largest cluster. Red, all other clusters. Black arrows, stress percolation events (red isolated filaments connect to the main cluster, becoming blue). Scale, 20μm/min **G:** PIV analysis of the bleb shown in panel 6E at times 0 and 35 s from the start of the acquisition. Squares, front and rear regions used in panels 6H and 6J. Colours, speed magnitude. Scale bar, 10μm. **H:** Mean PIV Y speed (velocity component parallel to bleb long axis) in the front and rear regions marked in panel 6G, as a function of time from the start of the acquisition (s, seconds). **I:** Segmented images of the bleb shown in panel 6E at time t = 0s and time t = 35 s. Blue, labelled C_1_, largest cluster. Red, all other clusters. L_C1_, maximum main cluster length. **J:** Renormalised maximum main cluster length L_C1_/L as a function of time from the start of the acquisition (s, seconds). Shadow, positions of the front (grey) and rear (cyan) regions from panel 6G-H. **K-M:** Left, renormalised maximum main cluster length L_C1_/L as a function of time from start of acquisition (s, seconds), for three representative bleb examples. Right, cluster analysis of the skeletonized actin network or one representative example. Blue, labelled C1, largest cluster. Red, all secondary clusters. **K:** Representative examples of persistent phenotype. **L:** Representative examples of intermittent phenotype. **M:** Representative examples of retracting phenotype. **N:** Boxplot of average occupancy 〈p〉 at the bleb front for persistent (white, P), intermittent (light grey, I) and retracting (R, dark grey) blebs. Dashed line, theoretical percolation threshold (p_c_ = 0.59). *P*, Welch’s T test.

### The advected percolation model predicts the various protrusion regimes observed experimentally

Some features of the steady-state regime of stable blebs, such as the actomyosin gradient, could also be predicted by the formerly proposed continuous fluid models (Callan-Jones and Voituriez, 2013; Ruprecht et al., 2015). We thus asked whether some of our observations that could not be explained by such models are predicted by the advected percolation model. This is the case for the pulsatile actin waves observed at the bleb front (Fig 6D-F). While they could not be predicted by the continuous fluid model (which is deterministic), pulsatile actin waves are predicted by the advected percolation model (which is stochastic). Experimentally, these fluctuations in the actin flow at the bleb front corresponded to phases of assembly of actin on the bare membrane front followed by phases of depletion, when the network reaches the percolation threshold and is connected to the flow produced by the contractile rear (Fig. 6E-F). On the model side, spatial fluctuations of the actin flow speed (Fig. 6G-H) can be explained by the fluctuations of the relative size *L*_*C*1_/*L* of the largest percolated cluster *C*_1_ (Fig. 6I-J). To test the model prediction, we plotted this model parameter (relative size *L*_*C*1_/*L* for the largest percolated cluster *C*_1_) for the different protrusion dynamics (Fig. 6K-M), which we divided into 3 categories: persistent, winding, and unstable (Fig. S4D-F and Movie S6). The model predicts that, as 〈*L*_*C*1_〉 ≈ *L_C_* approaches *L*, the forces are more likely transmitted to the front membrane and the bleb is thus more likely to retract or show unstable behaviours. Accordingly, we found experimentally that the average occupancy 〈*p*〉 at the bleb front is below *p_c_* (meaning *L_C_* is smaller than *L*) in persistent blebs, 〈*p*〉 » *p_c_* for retracting blebs, while in intermittent blebs 〈*p*〉 ≈ *p_c_* (Fig. 6N). Our modelling framework suggests that the various types of blebs described in the literature (“leader-bleb” (Logue et al., 2015), stable-bleb (Ruprecht et al., 2015), A2 migration (Liu et al., 2015), circus blebbing (Charras et al., 2008; Fujinami and Kageyama, 1975; Graziano et al., 2019), and bleb-on-bleb or sequential blebbing (Srivastava et al., 2020)), are emerging from the same simple advected percolation phenomenon and that they depend on the *L_C_/L* parameter. A recent independent study used similar theoretical tools to describe steady or pulsatile flow regimes in in vitro reconstituted actin networks (Krishna et al., 2022), showing the robustness of this approach. Overall, our advected percolation model, based on the basic ingredients gathered from the hNA TIRF observations, can explain complex aspects of bleb morphodynamics that could not be explained by the previous continuous fluid models of the actin cortex.

### Morphogenesis and motility of prototypic amoeboid motile cytoplasts are explained by the advected percolation model

Given the capacity of the advected percolation model to predict the actin structures and stability of stable blebs, we asked whether actomyosin follows the same organization in motile cytoplasts formed from stable blebs (Fig. 1G) and whether we can use our model to explain their motile behaviour and their shape. We recorded spinning-disc confocal timelapse images of motile cytoplasts, formed from actin-GFP expressing HeLa cells in a PLL-coated glass-bottom chamber (Fig. 7A-B and Movie S7). To perform a quantitative analysis of the relation between the actin cytoskeleton and the cytoplast shape and motility, we used a trained Ilastik model (Berg et al., 2019) and the Fiji-based plugin ADAPT (Barry et al., 2015) to obtain the outline of the cytoplast and compute the protrusion velocity (i.e. normal edge velocity) and the actin density along the cytoplast contour (Fig. S5B). This type of analysis has been performed before on lamellipodial migration (Ji et al., 2008; Ponti et al., 2004), but not on motile blebs. Plotting the average normal edge velocity for various levels of cortical actin density clearly showed that, contrary to lamellipodia, the protruding regions of the motile bleb cytoplasts were restricted to the lowest actin density regions, with a sharp threshold of density corresponding to a stalled edge and a negative edge velocity (retraction) for higher actin densities (Fig. 7C-E). This analysis thus identifies three regimes of membrane dynamics (protruding, stalled and retracting) which correspond to three ranges of actin concentration with sharp transitions, and three different shapes (the zero-velocity edge corresponding to the flat side, Fig. 7F). We propose that these ranges of actin concentration correspond to the three cortical states that we characterized using hNA TIRF on stable blebs (Fig. 7G), the percolated state corresponding to the stalled regime of membrane dynamics, while the contractile state would correspond to the retractile regime and the gas-like state to the protruding regime. This first analysis thus suggests that the locomotion and shape of motile cytoplasts might rely on the same actomyosin dynamics that we revealed using hNA TIRF microscopy in stable blebs.

**Figure 7:**
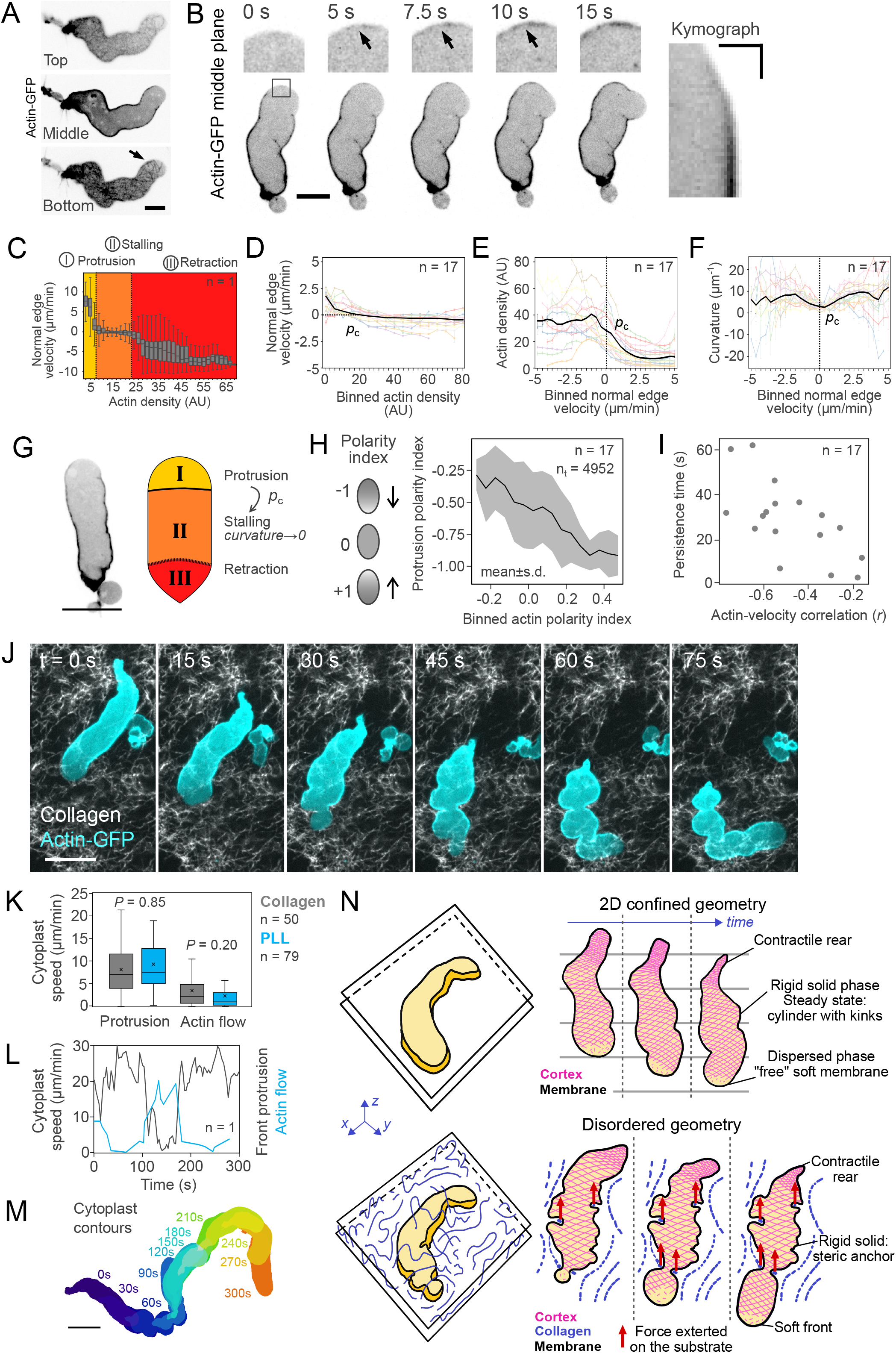
Cell confinement leads to the formation of motile cytoplasts. **A:** Representative fluorescence confocal spinning disk time-lapse images of a migrating cytoplast from 3μm-confined HeLa *ActB*-GFP cells (Actin-GFP) at bottom, middle and top planes. Arrow, sparse actin network at the front. Scale bar, 10 μm. **B:** Representative fluorescence confocal spinning disk time-lapse images of a migrating cytoplast from 3μm-confined HeLa *ActB*-GFP cells (Actin-GFP middle plane) under adhesive conditions (polylysine coating). Top, zoom on the front cortical region marked with a back square in bottom panel. Arrows, rebuilding of the cortical actin network. Time stamp, time elapsed from first frame in seconds. Bottom, full cytoplast. Scale bar, 10 μm. Right, kymograph of the front of the cytoplast shown in left panels. Scale bars, 5 μm, 5 s. **C:** Normal edge velocity as a function of the binned actin density of the cytoplast shown in panel 7A, calculated locally for 300 frames (150 s). Colours represent the three observed cortical regimes (yellow, protrusion; orange, stalling; red, retraction). **D:** Average normal edge velocity as a function of the binned actin density. Black solid line, average for all timepoints of all cytoplasts (n = 16). Coloured light lines, average for all timepoints for each individual cytoplast separately. Dashed black line, predicted percolation transition point, where velocity = 0. Bars, 95% confidence interval. **E:** Average actin density as a function of the binned normal edge velocity. Black solid line, average for all timepoints of all cytoplasts (n = 16). Coloured light lines, average for all timepoints for each individual cytoplast separately. Dashed black line, predicted percolation transition point, where velocity = 0. Bars, 95% confidence interval. **F:** Average curvature (μm^-1^, 1/radius) as a function of the binned normal edge velocity. Black solid line, average for all timepoints of all cytoplasts (n = 16). Coloured light lines, average for all timepoints for each individual cytoplast separately. Dashed black line, predicted percolation transition point, where velocity = 0. Bars, 95% confidence interval. **G:** Left, representative fluorescence confocal spinning disk image of a migrating cytoplast from 3μm-confined HeLa *ActB*-GFP cells (Actin-GFP) at middle plane. Scale bar, 10 μm.vRight, diagram representing the three observed cortical regimes (yellow, protrusion “I”; orange, stalling “II”; red, retraction “III”). The transition from the protrusion to the stalling regime occurs at the percolation density p_c_. The curvature at the stalling phase tends to 0. **H:** Left, diagram illustrating the polarity index. Accumulation at the rear yields a polarity index of +1, whereas homogeneous accumulation yields a polarity index of 0. Right, protrusion polarity index as a function of the binned actin polarity index (mean ± s.d., n = 17 cytoplasts and 4952 time points). **I:** Average persistence time (decay length of the edge velocity spatial autocorrelation averaged over all time points for a given fragment) as a function of the actin-velocity correlation (Pearson’s *r*). **J:** Representative fluorescence confocal spinning disk time-lapse images of a migrating cytoplast from 3μm-confined HeLa *ActB*-GFP (Actin-GFP, cyan) cells embedded in a collagen gel (Collagen, gray). Time stamp, time elapsed from first image in seconds. Scale bar, 10 μm. **K:** Average front protrusion (left, μm/min) and actin flow (right, μm/min) of cytoplasts migrating within a collagen get (grey, n = 50) or between two PLL-coated PDMS confinement slides (cyan, n = 79). Cross, mean values. Bars, median values. P values, Welch’s T test. **L:** Front protrusion (grey line) and retrograde actin flow (cyan line) as a function of time elapsed from first image in seconds, for a representative cytoplast shown in panel 7J. Note the slippage occurring between t = 100 s and t = 180 s. **M:** Colour-coded time projection of contours of a representative cytoplast shown in panel 7J. Time stamp, time elapsed from first image in seconds. Scale bar, 10 μm. **N:** Diagram of cortex properties during confined migration. Top left, schematic representing a confinement experiment in a free chamber. Top right, diagram representing an image sequence of confined migrating cell. Pink, mesh representing the actomyosin cortex. Bottom left, schematic representing a confinement experiment in a chamber filled with collagen. Blue, collagen fibres. Bottom right, diagram representing an image sequence of a confined migrating cell inside a chamber filled with collagen. Pink, mesh representing the actomyosin cortex. Blue, collagen fibres. Red arrows, forces exerted on the collagen gel by the migrating cell.

To examine more closely the morphodynamics of motile blebs, we performed a full characterization of the relation between the dynamic shape of the blebs and the density profile of the actin cortex, using cross-correlation analysis, similar to what was performed for mesenchymal protrusions (Ji et al., 2008). Because, similarly to stable blebs (Fig. S4D and Movie S6), motile cytoplasts displayed a variety of protrusion dynamics, we also classified them into 3 categories: persistent, winding, and unstable (Fig. S5B-C and Movie S6). Qualitative observation suggests that cytoplast shapes dynamics depends on actin density at the leading, forming persistent kinks which shape the cytoplast edges (Fig. S5B, black arrows and Movie S6). The spatiotemporal cross-correlation plots show the distinct modes of actin and membrane dynamics for each cytoplast category (Fig. S5C). Auto-correlation analysis of the edge velocity provides typical timescales and length scales for each cytoplast category, describing the size of the protruding region and its persistent time (Fig. S5D-E). Cross-correlation analysis combined with a close examination of the time-lapse movies provided additional insight into the spatiotemporal dynamics. It indicated, as expected from the simpler analysis above (Fig. 7C-F), that actin and edge velocity are negatively correlated with zero lag time (Fig. S5F) and that the rise in cortical actin precedes the membrane stalling (by 1.5 s), while membrane detachment (leading to protrusion) is followed by repopulation of the membrane with actin after a 9 s delay (Fig. S5G). More persistent migration was associated with lower actin-velocity correlation, confirming the relation between overall actin polarity and protrusion persistence (Fig. 7I). Taken together these results provide a quantitative description of the morphodynamics of the motile cytoplasts and confirm the main observation of an anti-correlation between actin density and protrusion speed, opposite to what has been described for lamellipodial motility (Ponti et al., 2004; Ryan et al., 2017).

According to our advected percolation model, the intermittent/winding/unstable regimes are due to transient events during which the main rigid/percolated actin cluster reaches the front edge (Fig 6K-N) and stalls or even retracts it, generating kinks (Fig. 5D). Reducing the strength of the actin/membrane binding should thus reduce the frequency of stalling and retraction events, leading to more stable blebs and more persistent motile cytoplasts. We found that pharmacological inhibition of the actin membrane linker ezrin using NSC668394 resulted in a decrease in the maximal force exerted by actin filaments on the plasma membrane in blebs (increase of snapping angle in Fig. S6A-B, increase of tearing angle in Fig. S6C-D) and cytoplasts (increase of tearing angle in Fig. S6E). Quantification of the occurrence of the three main categories of motile blebs showed that the persistent phenotype became largely dominant upon ezrin inhibition in both blebs and cytoplasts (Fig. S6F). Consistent with an increase in the fraction of persistent cytoplasts, Ezrin inhibition led to more actin accumulation at the back (higher actin polarity index, Fig. S6G), an increase in the slippage of the actin flow (Fig. S6H), and depletion of actin from the very front (Fig. S6I), but with a similar actin assembly rate (assessed by the slope of the actin intensity increase over time as it moves along the cytoplast edge, Fig. S6J). This effect of ezrin inhibition would correspond, in our model, to an increase in *v*_0_ with constant *β*, predicting a change in actin distribution and protrusion stability similar to what we observe experimentally. This increase in protrusion stability was reflected in an increase in edge velocity autocorrelation (a larger persistent width, Fig. S6K; and a longer persistent time, Fig. S6L). The change, compared to control, in actin-edge velocity correlation following ezrin inhibition (Fig. S6M-N) are coherent with a more polarized cytoskeleton and less cortical actin accumulation at protruding regions. Overall, these experiments confirm that, in motile blebs, the actin cortex attachment is antagonizing membrane protrusion. They also suggest that the morphodynamics of motile bleb cytoplasts can be qualitatively captured by our advected percolation model framework. Our model thus provides the right framework to capture cell scale actomyosin dynamics down to the single actin filament level in motile blebs. In conclusion, we believe that our observations and advected percolation model together draw a complete and consistent picture of an amoeboid-like motile cytoplast, providing an alternative locomotion mechanism, compared to the well understood lamellipodial driven mesenchymal motility (Theriot and Mitchison, 1991).

### Advected percolation explains efficient amoeboid migration through complex environments

We next asked whether bleb derived cytoplasts could migrate through complex environments such as collagen networks, thus recapitulating the capacity of amoeboid cells to efficiently navigate such environments. To test this hypothesis, we embedded TALENS-edited actin-GFP HeLa cells in a dense collagen matrix and deformed the gel until cells were confined to 3μm. We observed the formation of large blebs that did not retract (Fig. S5H arrows). Similar to the 2D pLL-coated condition (Fig. 1D), stable blebs migrated away from the cell while the cell body remained in place, and eventually produced motile cytoplasts. Cytoplasts tended to be of a larger size than stable blebs (Fig. S5I) and displayed a polarized actin cortex and seamlessly squeezed through small gaps, slightly deforming the surrounding collagen network (Fig. 7J and Movie S7). As a result, actin flow and protrusion speed were akin to the ones observed in PLL-coated 2D confiner devices (Fig 7K). In events where the cytoplasts migrated in a non-adhesive area, the protrusion speed decreased while the cortical flow slipped until a new grip was achieved (Fig. 7L-M), highlighting the role of cortical flow in driving cytoplast migration. Analysis of the collagen deformation field showed that motile cytoplasts produced expansile deformation at the leading edge and contractile deformation at the rear (Fig. S5J-L). Detailed observation of the actin network confirmed our working model: when the cytoplast faced a constriction, an actin free membrane protrusion first expanded through the gap, the actin cortex then densified at the gap region, likely corresponding to the rigid percolated network thus producing an anchor point, while the rear retracted, pushing the content of the cytoplast through the gap (Fig S5L). This series of events resembles what could be observed on motile cytoplasts and stable blebs in 2D confined spaces without collagen: the central advected rigid actin network would lead to the generation of irregularities on the bleb outline that then would travel from the front to the rear of the bleb with a fixed shape – in motile cytoplasts, these irregularities would travel from the front to the back of the cytoplast, while remaining stationary relative to the substrate. These observations suggest a way by which the percolating/solidifying actin network at the bleb front could shape itself to fit any external inhomogeneity of the environment and thus form anchors to which forces generated by the contractile rear can be transmitted to generate motion (Fig. S7N and Movie S7), even in the absence of friction forces. This could provide a complementary explanation to the mechanism by which environmental topography enables non-adhesive frictionless migration (Reversat et al., 2020). These observations confirm the unique locomotion capacity made possible by the advected percolated actin network in motile cytoplasts, with a soft actin-free front which can squeeze through small gaps, immediately followed by a percolated network forming stable anchors to the environment and finally a contractile rear producing the propulsion force and disassembling the network. Our work suggests that this seemingly complex series of actin structures can be explained simply with minimal ingredients and be produced spontaneously by actomyosin networks. We thus propose that the advected percolation phenomenon which we have described here both theoretically and experimentally constitutes a basic mechanism for efficient amoeboid locomotion through complex environments.

## Discussion

### A model for amoeboid cell motility down the single actin filament level and up to migration through complex networks

Our study of actomyosin dynamics in stable blebs using hNA TIRF microscopy allowed us to describe, at the single actin filament resolution, the dynamics of the network, and to understand its contribution to steady-state bleb shape (Fig. 5F). This description also proved valid to explain the motility of amoeboid bleb derived cytoplasts in 2D confinement and through collagen networks.

At the protrusion tip, actin filaments are sparse, short and not connected, which can be described as a gas-like phase. The front shape is that of a free membrane with uniform tension governed by Laplace’s law, and it can easily deform and insert through small gaps. In the middle region, the strain-free advected solid at steady state can only be cylindrical (to be translation invariant), giving the elongated shape and providing an anchor point in dense matrices. At the back, contractility leads to a convergent flow and thus a triangular shape, and generates rear retraction and actomyosin disassembly. We propose a physical model, which we call advected percolation, that demonstrates the capacity of actomyosin networks to spontaneously organise into the temporally stable spatial pattern of phases observed experimentally. The model is based on the combination of two basic ingredients: the occurrence of a rigidity percolation transition in the actin network at a critical density, and the advection of the percolated network powered by a self-sustained rear localization of myosin motors, which produces a gradient of filament density, thus maintaining the percolation transition away from the bleb front. Percolation in actin networks and its consequences on rheology (Alvarado et al., 2017, 2013), as well as contractile instabilities in actomyosin systems (Bendix et al., 2008; Callan-Jones and Voituriez, 2016; Mayer et al., 2010; Ruprecht et al., 2015), have been discussed before independently. Our study reveals the existence of cross-linked, percolated actin network, with a rigid solid-like rheology in the range of short time scales and low forces relevant to our observations. It occupies a large part of the motile bleb, between a loose network at the protruding front and a contractile network localized at the retracting rear. The function of this rigid network in leader-bleb shape and migration is consistent with the previous findings that actin crosslinkers are essential (Logue et al., 2015), since crosslinkers are essential elements to rigidify the network. Importantly, 3D imaging of the actin network in the stable blebs showed that they are hollow and that the actin network is forming a shell, justifying the use of a 2D percolation model (Fig. 7A). We argue that the combination of our detailed observations and our advected percolation model provides a complete picture for amoeboid migration and introduces a new mechanism of pattern formation potentially applicable to other synthetic or living active materials. The robustness of this mechanism is further supported by the independent study (Krishna et al., 2022), which points at similar self-organizing patterns in reconstituted actomyosin systems based on cell extracts encapsulated in water-in-oil emulsion and uses theoretical tools consistent with our description.

The molecular composition and dynamics of actin structures have been described in detail in many biological systems, in which they often produce forces to drive morphogenesis and motion (Heisenberg and Bellaïche, 2013; Svitkina, 2018). One of the first simple actin-based migration machineries understood from first principles down to single actin filaments was the propulsion by actin comet tails of intracellular bacteria such as *Listeria monocytogenes* (Loisel et al., 1999; Theriot et al., 1992). The understanding of actin comet tails allowed to propose simple models for the formation of various membrane-enclosed cellular protrusions such as filopodia and, together with a wealth of additional studies (Keren et al., 2008; Svitkina and Borisy, 1999; Watanabe and Mitchison, 2002), eventually allowed to draw a clear picture of lamellipodial protrusion driven by actin polymerization. Several studies have also described the formation of the actin cortex in transient, retractile blebs (Charras et al., 2006; Chikina et al., 2019). In this work, we complete this picture of actin-driven locomotion by explaining amoeboid motility, from first principles down to single actin filaments.

### From transient to stable blebs

Blebs, which are characteristic of amoeboid migration, have been mostly described as transient protrusions which retract within minutes. Nevertheless, the blebs observed here can be stable for several hours. The advected percolation model we propose above is meant to explain the steady-state regimes, it can however also make simple qualitative predictions on the initial bleb dynamics (Fig S1). In brief, if actin assembly starts in the absence of myosin, and therefore in the absence of advection, it will lead to a uniform increase of actin filament concentration, which will eventually uniformly reach percolation, filling the entire bleb. Myosin action on such a network will retract the fully connected (solid) bleb, producing a transient bleb. On the contrary, if myosin activity builds up fast enough, a gradient of actin filaments can be formed, leading to a stable bleb with *L_C_* < *L* (*v*_0_ large in the model). To test this prediction, we focused on the early time points after confinement. Upon confinement, cells initially formed round transient blebs, while long and more stable blebs were visible after a few tens of seconds, when myosin II is activated (Fig. 1A and S1). As reported in other works (Cunningham, 1995), transient blebs inflated to a maximum size within a few seconds, reassembled a cortex, and completed retraction within a minute (Fig. S7A-C, transient bleb). On the other hand, stable blebs reached a stable elongated shape where actomyosin was depleted from the leading edge and the projected area remained stable (Fig. S7A-D, stable bleb). We systematically quantified the three parameters that separate stable from transient blebs: their lifetime, shape, and polarity of actomyosin. We plotted, for every single bleb, its aspect ratio versus its polarity index and colour-coded its lifetime. This plot shows two well-defined groups of blebs: round, homogeneous, short-lived blebs versus elongated, polarized, and long-lived blebs (Fig. S7E). The lack of intermediate phenotypes, which is even more evident on the NMIIA polarity graph, suggests that, while all blebs initiate from round actin-free membrane protrusions, their evolution in time diverges into two groups, one which forms a front cortex and retracts, and one which forms a gradient of actomyosin and remains stable, suggesting a sharp transition between the two states, as qualitatively predicted by the advected percolation model. To quantitatively analyse the temporal patterns across many stable and transient blebs, we aligned and averaged the projected area dynamics, the front actin density, and the coefficient of variation of the NMIIA signal relative to the flow onset (Fig. S7F and Movie S1, stable bleb part). Consistent with our hypothesis, stable blebs never displayed actin accumulation at the front tip, whereas transient blebs built a front cortex ~15 s before flow onset (Fig. S7F and Movie S1, transient bleb). We plotted the relative time of maximal membrane extension (membrane stalling) and flow onset after bleb initiation, colour-coding bleb stability. The time of flow onset in transient blebs was always longer than the time of membrane stalling, and the opposite was true for stable blebs (Fig. S7G). The difference between the timing of actin cortex assembly and the onset of NMIIA contractile activity thus determines whether the retrograde flow starts before or after the actin cortex has populated the entire bleb membrane and, in turn, determines the fate of the bleb. In other words, the reach of the percolating cluster relative to the total bleb size determined the stability of the bleb upon contraction onset (Fig. S7H-I). In conclusion, our model and observations provide a simple explanation for the sharp transition between transient and stable blebs upon activation of NMIIA activity right after cell confinement.

### Physiological relevance of stable blebs and motile fragments

Our observation and model can thus explain the motility of motile bleb cytoplasts through complex networks. The formation of large long-lived or stable blebs has been reported in a variety of cell types during *in vitro* confinement. These structures can drive the migration of the entire cell and might help cancer cells squeeze through the matrix in an amoeboid fashion during the metastatic process. In addition, migrating cells have been shown to form motile cell cytoplasts with different biological functions (e.g. horizontal transfer of bioactive material, migration guidance), called cytokineplasts or motile cytoplasts. Early studies documented the formation of cytokineplasts from detaching cell protrusions *in vitro*, but a recent study observed, in human xenograft tumours *in vivo* in mice, that motile cytoplasts were formed from large blebs (Headley et al., 2016). These cytoplasts migrated away from the tumour autonomously for hours despite the absence of a nucleus, eventually modulating the interaction between the immune cells and the tumour. Our work might explain the formation and persistent motility of these cytoplasts.

Transient blebs have been implicated in particular forms of amoeboid migration, called bleb-based migration, especially in cancer cells. In this mode of migration, transient blebs produce either pushing or pulling forces due to their fast cycles of extension and retraction (Charras and Paluch, 2008; Welf et al., 2020a). The mechanism we propose here is different, as it produces a stable polarised migrating object, rather resembling fast migrating cells such as confined neutrophils (Graziano et al., 2019). This type of migration has now been described in a large variety of cell types and organisms and has been proposed to represent an ancestral conserved migration machinery (Fritz-Laylin, 2020; Liu et al., 2015). Beyond amoeboid bleb-based migration and locomotion of cell-derived cytoplasts, recent work pointed to the possible role of bleb-like membrane detachment even in mesenchymal lamellipodial driven migration, the formation of protrusions being always preceded by the depletion of ERM proteins (Welf et al., 2020b). These observations led the authors to propose an alternative model for the initiation of protrusions, in which actin mostly holds the membrane and hydrostatic pressure rather than actin pushes the membrane. This vision is consistent with another recent study (Bisaria et al., 2020) which demonstrated that the actin network in close contact with the membrane hinders membrane protrusion. Membrane-proximal actin is depleted in front protrusions and enriched at the cell rear, forming a gradient at the scale of the entire cell, similar to what we observed in the simpler context of motile bleb cytoplasts in which other actin structures are not present. Together, these studies suggest that our advected percolation model might be relevant not only for bleb-like protrusions and amoeboid cells, but also for mesenchymal migration, providing a simple generic principle for cell locomotion.

Our advected percolation model seems sufficient to explain the main features of a minimal motile system. We anticipate that this model might open the way to the reconstitution from purified components of cell locomotion. Nevertheless, because of the advection of the percolated network, in the molecularly complex context of the cell, a large number of other constituents might also be advected and thus spatially organized, maybe contributing to the general stability of the bleb organization and the persistence of the locomotion. Further work will be needed to strictly determine the minimal ingredients necessary to reconstitute an autonomous motile cell-like system. Beyond the reconstitution of cell locomotion from purified components, our advected percolation model could also inspire the engineering of innovative materials and robots (Hawkes et al., 2017; Kim et al., 2013) based on its simple principle – a soft front solidifying and being then disassembled by a force producing rear, leading to autonomous machines capable of efficient locomotion through complex environments.

### Limitations of the study

Our study is fully based on detailed recording and analysis of the motion and shape of the stable blebs and the underlying actin network, but we do not provide any measure of the forces involved in these processes. The forces involved in cell migration have been investigated in several previous studies using traction force microscopy (Reinhart-King et al., 2003). Nevertheless, these forces are accessible mostly in the context of mesenchymal, adhesion-based migration, and are associated with the formation of strongly adhesive structures and associated stress fibres, which are not present in fast amoeboid migration. It has proven difficult to measure these forces for amoeboid cells, as reported in previous studies (Bergert et al., 2015; Reversat et al., 2020), mostly because the forces are too small and there is currently no reliable technique available to measure them. Similarly, forces within the actin network would be important to determine, but this has never been achieved so far in a living cell. Our assessment of whether the network is rather solid or fluid is mostly based on a geometric definition (whether the filaments move relative to each other in the network). It is likely that under sufficient forces, the percolated network might display classical visco-elastic properties. Nevertheless, under the conditions studied here, the timescale and the forces involved in this type of migration, the percolated network behaves geometrically like an advected solid.

## Materials and methods

### RESOURCE AVAILABILITY

#### Lead contact

Further information and requests for resources and reagents should be directed to and will be fulfilled by the lead contact, Matthieu Piel.

#### Materials availability

This study did not generate new unique reagents.

#### Data and code availability

All original microscopy images and datasets shown in the figures have been deposited at Figshare and are publicly available as of the date of publication. DOIs are listed in the key resources table. Additional microscopy data reported in this paper will be shared by the lead contact upon request.

All original code and associated datasets have been deposited at Figshare and are publicly available as of the date of publication. DOIs are listed in the key resources table.

Any additional information required to reanalyze the data reported in this paper is available from the lead contact upon request.

### EXPERIMENTAL MODEL AND SUBJECT DETAILS

#### Human cell lines

Human cervical adenocarcinoma cells HeLa-Kyoto stably expressing myosin IIA (Myh9)-GFP and LifeAct-mCherry or Myh9-GFP, or LifeAct-mCherry and the plasma membrane-targeting CAAX box fused to GFP, or TALEN-edited ActB fused with GFP (Cellectis, Paris, France), or with no stable marker were cultured in DMEM GlutaMAX medium (Gibco, #61965-026) supplemented with 10% FBS (Gibco, #P30-193306A) and Penicillin Streptomycin (Thermofisher, #15140-122) at 37°C and 5% CO2.

All cell lines were tested for mycoplasma contamination using Mycoplasmacheck PCR Detection (Eurofins, #50400400), and treated once a year with Plasmocin profilactic (InvivoGen, #ant-mpp) and treatment Mycoplasma Elimination Reagents (InvivoGen, #ant-mpt).

### METHOD DETAILS

#### Construct and DNA transfections

Cells were transfected with plasmid DNA using Lipofectamine 3000 reagent (Thermofisher #L3000001) transiently or stably, according to manufacturer’s protocol, and imaged from 24 to 72 hours after transfection. The following expression vectors were used for plasmid DNA transfections: pCMV-mEmerald-FilaminA-N-9 (Addgene #54098, gift from Michael Davidson), pCMV-mCherry-FilaminA-N-9 (Addgene #55047, gift from Michael Davidson), pCMV-mEmerald-Alpha-Actinin-19 (Addgene #53989, gift from Michael Davidson), pCMV-mCherry-Alpha-Actinin-19 (Addgene #54975, gift from Michael Davidson), pCMV-mApple-MyosinIIA-N-18 (Addgene #54930, gift from Michael Davidson), and pEGFP internal GFP Ezrin (gift from Florence Niedergang).

#### Drug treatments

The following pharmacological inhibitors and chemical compounds were used: 10 μM ezrin inhibitor NSC668394 (EMD Millipore #341216), 10 μM ROCK-mediated contractility inhibitor Y-27632 (ED Millipore #688000), 20 μM AACOCF3 inhibiting the nuclear envelope stretch-sensitive enzyme cPLA2 (Tocris Bioscience #1462), 100 μM 2-APB blocking stretch-activated inositol triphosphate receptors (InsP3Rs) on the ER/nuclear membranes (Tocris Bioscience #1224), 100 μM CK-666 (Tocris Bioscience #3950) inhibiting Arp2/3 inhibiting actin polymerization, and 40 μM SMIFH2 inhibiting FH2 domain inhibitor preventing actin nucleation (Tocris Bioscience #4401)

Growth medium was supplemented with 1% DMSO (vol/vol) (Sigma-Aldrich #D2438) in control experiments when DMSO was used as a solvent. Cells were imaged 30 minutes after treatment. For AACOCF3 treatment, medium in perturbation and control experiment did not contain FBS. The PDMS devices in contact with cells during imaging were incubated in medium supplemented with the final concentration of drug during at least 1h before the experiment.

#### Fabrication of the confiner PDMS device

We used a single-well confiner device developed in our lab for cell confinement, which combines confinement precision and high imaging quality (Le Berre et al., 2014). The device consists of a pressure cup, a confiner coverslip, and a glass-bottom dish. The pressure cup was made with a custom-made metallic mould from polydimethylsiloxane (PDMS; Neyco, #RTV615) at 10:1 proportion (10/1 w/w PDMS A / crosslinker B). Confinement was controlled by a Flow EZ pressure system (Fluigent, #LU-FEZ-N800, #LU-SPK-0001, #LU-LNK-0001, #SFT-MAT, #32305001). To make the confiner coverslip at the desired height, 12 mm glass coverslips were plasma treated and then placed on top of a PDMS mixture 10:1 on the wafer moulds (containing holes of 3 μm depth, fabricated following standard photolithography procedures). The height of the micropillars determines the height for spatial confinement of cells between the coverslip and the substrate, which is 3 μm. After baking at 95°C for 15 min, coverslips with PDMS pillars were carefully removed from the wafers under isopropanol, cleaned with isopropanol, and air-dried. For non-friction confinement conditions, the glassbottom dish (World Precision Instruments #FD35) and the microfabricated confiner coverslips were treated with plasma for 1 min, and incubated with 0.5 mg/mL pLL-*g*-PEG (SuSoS, PLL(20)-g[3.5]-PEG(2)) in 10 mM pH 7.4 HEPES buffer for 1h at room temperature. For friction confinement conditions, the device was instead coated with a 0.5 mg/mL of PLL solution in PBS diluted from Poly-L-lysine solution 0.1 % (w/v) (Sigma # P8920). Confiner coverslips and glass-bottom dishes were rinsed and incubated in medium for at least 2 hours before confining the cells.

#### PDMS device confinement and live imaging

Trypsinized (Gibco # 10043382) cells were resuspended in culture medium and plated on the device just before applying the confinement. In the case of high NA TIRF imaging, the setup does not have CO2 regulation, so cells were resuspended instead on CO2-independent medium (Gibco, #18045088).

Confocal fluorescence time-lapse recordings were acquired with 63x oil (HC PL APO 63x CS2, NA: 1.40, #11506350) or 100x oil (HC PL APO 63x CS2, NA: 1.40, #11506372) objectives with a Yokogawa CSU-X1 spinning-disc head on a DMI-8 Leica inverted microscope equipped with a Hamamatsu OrcaFlash 4.0 Camera, a NanoScanZ piezo focusing stage (Prior Scientific) and a motorized scanning stage (Marzhauser), controlled by MetaMorph software (Molecular Devices). The spinning disk confocal microscope was equipped with an on-stage incubation chamber which maintained the temperature at 37°C and CO2 concentration at 5%.

The high NA TIRF microscope consisted of a house-made TIRF setup with standard settings, equipped with a 473nm laser (gem473 500mW, LaserQuantum), a 100x TIRF objective (Olympus, NA: 1.49, UAPON 100XOTIRF), and a sCMOS camera Andor Zyla 4.2 (Oxford Instruments). For two-colour TIRF, an additional 561nm laser (gem561 500mW, LaserQuantum) was used, and the acquisition was performed sequentially. A custom-made FPGA trigger was used for the lasers, synchronized with the camera. A single notch filter was used in the emission light path to block the laser lines 405/473/561 (Chroma). Acquisition was controlled by the Andor SOLIS software (Oxford Instruments).

#### AFM confinement and live imaging

Trypsinized cells were suspended in CO2-independent, phenol red-free DMEM/F-12 medium supplemented with 10% FBS (Invitrogen) and plated on PLL-*g*-PEG-treated glass-bottom 35-mm dishes (FluoroDish, WPI). These were mounted in a dish heater (JPK Instruments) and kept at 37°C under an inverted light microscope (Axio Observer.Z1; Zeiss) equipped with a confocal microscope unit (LSM 700; Zeiss) and the atomic force microscope (AFM) head (CellHesion 200; JPK Instruments). Experiments were initiated 30 minutes after cell plating to allow for cell sedimentation.

Focused ion beam (FIB)-sculpted, flat silicon microcantilevers were processed and calibrated as described to confine cells in a parallel plate assay (Cattin et al., 2015). The microcantilever was fixed on a standard JPK glass block and mounted in the AFM head. The cantilever was lowered at constant speed of 0.5 μm·s^-1^ on the cell until reaching a pre-set height, and the resulting varying cantilever deflection (force) were recorded over time. At the same time, differential interference contrast (DIC) and fluorescence images at the confined cell midplane were recorded every 5 s using a 63× water immersion objective. Imaging and measurements were carried out in a custom-made isolation box. All AFM experiments were done in cell culture medium as specified.

#### Collagen-embedded confinement

Confinement experiments with cells embedded in collagen were conducted as follows: i) collagen gel preparation at 4°C to avoid polymerization; ii) cell trypsinization, washing, and mixing with the collagen solution; iii) confinement to 20 μm and 37°C during 30-60 minutes to allow collagen polymerization; iv) confinement to 3 μm to allow stable bleb formation; v) imaging of cytoplasts migrating through the matrix. The gel was prepared using the following reagents (per ml of solution): 100 μl MEM10x (Thermofisher #11430030), 50 μl NaOH 0.34N (Sigma-Aldrich #S2770), 25 μl HEPES 1M (Sigma-Aldrich #H0887), 25 μl of cy5-labelled type-I collagen solution (Jena Bioscience #FP-201-CY5) and completed with water and Collagen type-I rat tail solution (Corning #354236). The confiner device was treated with 0.5 mg/mL pLL-*g*-PEG before the experiment.

### QUANTIFICATION AND STATISTICAL ANALYSIS

#### Statistical analysis of experimental data

Statistical analyses for all experiments were performed in Python 3 (NumPy and SciPy libraries) or Microsoft Excel. Statistical data are presented as median or mean ± standard deviation or interquartile range. For each panel, sample size (*n*), experiment count (*N*) statistical tests used, and *P* values are specified in the figure legends. Samples in most cases were defined as the number of cells examined within multiple different fields of view on different dishes, and thus represent data from a single experiment that are representative of at least three additional independently conducted experiments.

#### Processing and quantifications of experimental data

All image acquisition data were processed using Fiji (NIH).

#### Analysis of bleb and actomyosin dynamics in confocal images

Confocal fluorescence images were background subtracted and bleach corrected using Fiji (https://imagej.net/Bleach_Correction).

*Time of bleb nucleation* (relative to the beginning of compression), *bleb lifetime* (time between bleb nucleation and full retraction), and *bleb aspect ratio* (bleb length divided by bleb maximal width) were measured using the custom-made Fiji macro “bleb_morphogenesis.ijm”, which performs radial reslicing and facilitates manual tracking.

*Cortical actomyosin density* was calculated using a manually-selected 500nm-wide region on images with subtracted cytoplasmic background In time analysis, these regions were manually selected at every time point. To compare cortical gradients between round and elongated blebs, the length was normalized by dividing it by the total, and mean intensities were calculated as a function of bleb position divided into 10 bins: (0,0.1], (0.1,0.2],… (0.9,1.0], taking 0 for the bleb base and 1 for the bleb front

*Bleb or cytoplast projected areas* were calculated in Fiji from manually selected regions.

*Coefficient of variation of the fluorescence signal* was defined as *CV* = *σ/μ*. *CV* was calculated from images with the subtracted minimum cytoplasmic signal. Regions were manually segmented and signal standard deviation and mean were calculated using Fiji.

*Steady-state actin flow* and *protrusion* velocity were measured from kymographs on the myosin or actin channels taken at the bottom plane of cells, blebs, or cytoplasts. The instantaneous flow or protrusion speed at time *t* was defined as the mean slope in the kymograph around time *t*, manually measured. In cases where the flow was measured at each timepoint, *particle image velocimetry* (PIV) analysis was used instead, as described below.

To calculate the *actin polymerization dynamics* in the cortex, we first subtracted the background inside the cell, outside of the actin shell. Then, we manually selected 2 by 0.5 μm windows on the cortex as they moved backwards with the retrograde flow and calculated the intensity at each time frame. The curves were divided by their mean value on the plateau reached after 10-20 s. The slopes were then calculated, doing a linear fitting on the range (0.25-0.9).

*Polarity index* in blebs was defined as PI = (ρ_real_-ρ_front_)/(ρ_rear_+ρ_front_), where *ρ_rear_* is the integrated actin/NMIIA intensity in the cell-proximal or rear half of the bleb or cytoplast, and *ρ_front_* the integrated actin/NMIIA intensity on the other half. The estimated cytoplastic background is subtracted before calculation. In cytoplasts, *polarity index* was instead calculated from 500-nm wide regions along the cytoplast contour. Cytoplast contour was segmented with a user-trained Ilastik model (Berg et al., 2019). The segmented images and the actin-GFP channel were processed using the Fiji plugin ADAPT (Automated Detection and Analysis of ProTrusions) (Barry et al., 2015). The parameters used were as follows: smoothing filter radius: 1, erosion iterations: 2, spatial filter radius (μm): 0.5, temporal filter radius (s): 1, cortex depth (μm): 0.5, Visualization line thickness: 2. The output of ADAPT is an edge velocity and actin density map (summed over the 0.5 μm-wide cortical region). To detect the protruding front, ADAPT output was processed using the custom-made Python script “ADAPT_processing.ipynb”, which also includes the cross-correlation analysis described below.

*Distal actin density* was calculated manually selecting cortical hemispherical regions at the bleb front, measuring the actin density, and subtracting the mean cytoplasmic background.

*Cortical flow onset* was tracked visually assessing the start of contraction at the bleb rear.

*Maximal area timing* was defined as the first timepoint where the bleb projected area stopped to grow, assessed from manually selected regions on Fiji.

*Mean collagen deflection* map was calculated using the collagen fluorescence channel during and after cytoplast migration. These pairs of collagen images were compared using *particle image velocimetry* (PIV) analysis on the custom-made plugin “collagen_deflection_PIV.ijm”. PIV vectors from different fragments were averaged using the custom-made Python script “collagen_deflection_mean.ipynb”

#### Analysis of AFM confinement experiments

The raw AFM force data contains cantilever height, cantilever deflection (force) and time. The custom-made Python script “AFM_data_processing.ipynb”: i) identifies the AFM calibration steps to calculate the precise confinement height; ii) identifies the beginning of confinement, defined as the first time point at which the target cantilever height is reached (in this case 3μm); and iii) aligns the force curves by the time of the beginning of the confinement. The script also identifies the time of active force onset. To do so, force curves are first applied a rolling mean over 5 s. Then, the force onset timepoint (change of trend) is defined as the first time point where d*F*(*t*)/d*t* = 0, where *F*(*t*) is the AFM force.

*Mean edge retraction speed* was calculated on the segmented actin channel using the Fiji plugin ADAPT “Automated Detection and Analysis of ProTrusions (Barry et al., 2015).

*Number of blebs per cell* was calculated defining four random sites equally spaced in the cell and measuring the time of appearance, length, and width of all blebs in these sites during 300 s following confinement.

#### Analysis of single actin filaments

*Actin organization* and single filaments in blebs in Figure 2 were segmented and tracked manually using Fiji. Filament position was used to calculate *rotational velocity, mean square angular displacement, mean square displacement, growth speed*, and *branching angle*. Single filament growth analysis included only nascent branches identifiable with more than 15 consecutive frames in focus and isolated.

*Rotational velocity* was calculated as v_rot_ = Δθ/Δt = (θ(t_1_)-θ(t_0_))/(t_1_-t_0_).

*Mean square displacement* was defined as 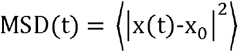 and the *mean square angular displacement* as 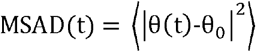.

*Nucleation of network segments* by branching or seeding was tracked manually using Fiji.

#### Analysis of cortical flow

*Particle image velocimetry* (PIV) analysis on high numerical aperture TIRF images was performed in Fiji using the custom-made plugin “PIV_analysis-stack.ijm”. Before analysis, TIRF images were normalized by dividing each time frame by its mean intensity value and subtracting the cytoplasmic background. The “PIV_analysis-stack.ijm” macro uses the “iterative PIV” plugin (available at https://sites.google.com/site/qingzongtseng/piv) initially developed for force traction microscopy, removing in the parameters any vector interpolation bias (Tseng, 2011; Tseng et al., 2012). The “iterative PIV” plugin was primarily based on JPIV, an open-source PIV software, which used standard algorithms to calculate the displacement vectors (available at https://eguvep.github.io/jpiv/). The interrogation window was estimated to be large enough from the images’ feature size, and the search window was estimated to be large enough to capture the maximal displacement observed in the images, such as search window size = interrogation window size + 2 × maximal displacement. The *x, y, ux, uy* values were imported into the custom-made Python script “cortical_flow_analysis.ipynb” script for processing and data analysis.

*Cortical flow divergence* and *net network turnover* were calculated using the custom-made Python script “cortical_flow_analysis.ipynb”. The divergence was calculated as 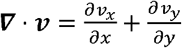. The divergence of the velocity field is a good indication of the relaxation or contraction of the cortex. Knowing the density and the cortical flow, one can calculate the flow divergence and the actin turnover rate. The spatial distribution of network turnover in 2D can be calculated from the mass conservation equation 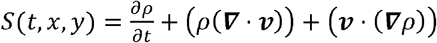, where *ρ* is the fluorescence intensity and *S* the network turnover (Yolland et al., 2019). We will consider steady states, so 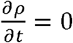, and we can drop the time dependence in *S*. This implies that *S*(*x, y*) = (*ρ*(*∇* · *v*)) + (*v* · (*ρv*)) = *∇* · (*ρv*), where the last equality is a mathematical identity. Therefore, knowing *v*(*x, y*) and *ρ*(*x, y*) one can calculate the local network turnover.

*Local alignment c*(*r*_0_) was defined as the average cosine similarity 〈cosθ〉 between all velocity vectors and their 8 nearest neighbours, and was computed using cos*θ* = *v*_1_ · *v*_2_/ ||*v*_1_||||*v*_2_|| (Yolland et al., 2019). For simplicity, we selected only the y-component of the velocity vectors. This algorithm was implemented in the “cortical_flow_analysis.ipynb” script and the averaged *C*(*r*_0_) over time was calculated for each window position.

*Velocity spatial correlation function* was defined as: 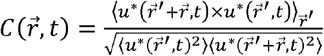, where 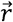 is the correlation distance and *u*\* is the normalized velocity such as *u*\* = *u* – 〈*u*〉 (Petitjean et al., 2010). This function averages across many pairs of points at distance 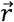 at a specific time. Using *u*\* instead of *u* gives the good normalization, such as 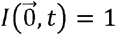 for all times and 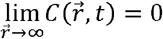. We take an averaged correlation over all directions, so 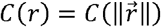, define the centre cell as (*x, y*) = (0,0) with an autocorrelation value *C*(0) = 1, and define *u*\*(0) as the velocity of the cell in the centre, our reference to calculate the correlation. The correlation at distance *r* from the centre *u*\*(0) will be: 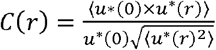, which simplifies to 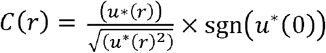. This satisfies the condition *C*(0) = 1, as 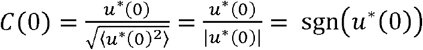. For our purposes, we defined the correlation distance *C*(*r*) as the closest distance at which *C*(*r*) < 0.5. To simplify the analysis, we took only the Y component of the velocity vector, as the x-component does not represent a significant part of the velocity. This algorithm was implemented in a the “cortical_flow_analysis.ipynb” script and the averaged *C*(*r*) over time was calculated for each window position.

*Bleb contour curvature* |κ| was calculated on the Fiji plugin “Kappa”, based on manually-drawn bleb contours. Curvature was taken in absolute values.

Tracking of landmarks (polygon vertices) in the flowing actin network was done by tracking manually frame by frame cortex structures and features.

#### Analysis of percolation in the actin network

The central concept of this work is *percolation*. Percolation theory describes the behavior of a network as a function of the parameter p, which controls the *occupancy* of sites or bonds of a lattice, thus relating microscopic (e.g.: the density of network components 〈p〉 and macroscopic features. Percolation models can predict geometric phase transitions, since at the critical point p_c_ the network suddenly changes its properties (Stauffer et al., 1994). That is, the percolation probability ⊓ (the probability of finding a path in the network that crosses the entire lattice) suddenly increases at the critical point p_c_. Theoretical or simulation based calculations predict that the value of p_c_ changes for different lattice types or if site or bond percolation are considered (Stauffer et al., 1994).

For ⊓, the network is composed of small disconnected clusters. Around the critical point 〈p〉≈p_c_, the network displays a broad distribution of sizes of clusters, with in particular the emergence of a cluster whose size is comparable to the entire lattice. For 〈p〉>p_c_, the network contains one large cluster, so-called spanning cluster. Therefore, the size of largest C_1_ and second largest C_2_ clusters evolve differently at *p_c_*. The size of C_1_ increases monotonically to reach a maximum after *p_c_*. In contrast, the size of *C*_2_ reaches a maximum at *p_c_*, where C_1_ ≈ C_2_, to then decrease at 〈p〉>p_c_ (Margolina et al., 1982). At the critical point 〈p〉≈p_c_, many network properties behave as of a power-law with |p-p_c_|. Scaling theory predicts the existence of universal critical exponents, which characterize these power-law relationships. The Fisher exponent ⊓ characterizes the distribution n_s_ of clusters of size *s*, such that n_s_~s^-τ^ for ⊓. The Fisher exponent is estimated to be τ≈2 for most systems, so that n_s_~s^-2^. The exponent ⊓ characterizes the divergence of the correlation length ⊓ around the percolation threshold, so that *ξ*~|*p*-*p_c_*|^-v^. The correlation length is defined as the distance over which a specific thermodynamic variable in the system are correlated with one another. In our case, for the flowing actin cortex, we measure the spatial velocity-velocity correlations C(r), which is also a measure of the linear extent of the spanning cluster. At 〈p〉>p_c_, the spanning cluster extends over the whole system, and the system becomes homogeneous (Alvarado et al., 2017; Stauffer et al., 1994).

Classical percolation theory analysis assumes sufficiently large length scales, as well as spatially homogeneous p. In an infinite system (⊓), the percolation probability ⊓ is ⊓ = l above *p_c_* and ⊓ = 0 below *p_c_*. However, for finite systems with large *L*, ⊓ can be written as a scaling function *Φ* ranging from 0 to 1. Practically, this means that the ⊓ function goes from a step function to a sigmoid-like function with decreasing system size (Stauffer et al., 1994). Likewise, percolation critical exponents are corrected for finite-size system. For example, if the correlation length approaches the size of the system, power-law scaling is no longer applicable. In our case, we test qualitatively general assumptions from percolation theory and complete this description with an analytical model and simulations. The consequences of flow, gradients, and finite system size to percolation critical exponents and network behavior are not fully understood and remain an active field of research. Moreover, a precise powerlaw fitting at such small scales is inconclusive due to large finite-size effects.

To segment the actin filaments from cytoplasmic background in the TIRF movies, we trained an Ilastik model (Berg et al., 2019). This allowed us to classify the image pixels in actin versus background pixels. The *occupancy* parameter p of a region was defined as the fraction of actin pixels over the total pixels, 〈*p*〉.

To analyze the *percolation probability* ⊓ as a function of the *occupancy p*, we divided the bleb in small windows along the longitudinal axis. Because the *occupancy p* is a function of the bleb length, this allowed us to compute the *percolation probability* ⊓ without a large gradient. For every single window, the *percolation probability* was set to ⊓ = 1 if a connection path was found across the window and ⊓ = 0 if no connection path was found. The average *occupancy* 〈*p*〉 was calculated for every window. Finally, the average *percolation probability* 〈⊓〉 was plotted as a function of the binned average *occupancy* 〈*p*〉.

*Cluster sizes* were defined in individual windows as the width of clusters normalized by the total bleb width. For the analysis plotting the cluster sizes as a function of the binned occupancy, all bleb windows were pulled together.

*Cluster areas* were defined in individual windows as the convex area of clusters. For the analysis plotting the cluster size distribution as a function of binned occupancy in log-log space, all bleb windows were pulled together.

*Largest cluster extension* ⊓ was defined as the size of the largest cluster over the bleb longitudinal axis, normalized by the bleb size.

The *average occupancy at the bleb front* ⊓ was calculated fitting a hemispherical region at the bleb front, and then measuring the average *occupancy* 〈*p*〉.

All the percolation analysis was implemented in two subsequent Fiji macro and Python script. Images were processed using the custom-made Fiji plugin “percolation_processing.ijm”, which outputs data tables. These tables and the PIV data from “cortical_flow_analysis.ipynb” are used as a input for the custom-made Python script “percolation_analysis.ipynb”, which outputs the plots.

#### Analysis of membrane dynamics and blebbing regimes

*Tearing angle* was defined as the angle between the tangent of the bleb membrane and the tangent of the cell body, and manually tracked using Fiji as previously (Charras et al., 2008). Tearing is theoretically expected to occur if f = T_m_(1-cos(180-θ))>J, with *θ* being the outer angle between the bleb membrane and the cell body and *J* being the adhesion energy between the membrane and the cortex. Therefore a larger *θ* implies higher *T_m_* or lower *J*.

*Snapping angle* was defined as the maximum indentation that single actin bundles could exert on the bleb front, and was manually tracked using Fiji.

*Blebbing regimes* were assessed based on actin and membrane dynamics at the bleb front. Persistent blebs were defined as bleb with no actin scars and persistent tip, without significant perturbations in the blebbing angle. Winding blebs were defined as blebs with alternating actin scars on the middle plane in the actin channel, and with a winding membrane tearing at the bleb front. Unstable blebs were defined as blebs that keep a constant blebbing at the cell front while transiently rebuilding a cortex all along the cell contour. To calculate the *fraction of blebs* in a given regime in a given condition, we recorded time-lapse movies on the actin and brightfield channels. All stable blebs present in each field of view were classified every minute based on the characteristics of the actin cortex and membrane dynamics, and the data pooled together. To classify the regime of cytoplasts dynamically, movies were divided in 25 s intervals and blebs were manually assigned a dominant behaviour in each time window.

#### Cell outline tracking and cross-correlation analysis

To extract the spatiotemporal dynamics of actin cortical density and protrusion speed, middle plane movies of actin-GFP cytoplasts taken on a confocal spinning disk were segmented using a user-trained Ilastic model (Berg et al., 2019). The segmented images and the actin-GFP channel were processed using the Fiji plugin ADAPT (Automated Detection and Analysis of ProTrusions) (Barry et al., 2015). The parameters used were as follows: smoothing filter radius: 1, erosion iterations: 2, spatial filter radius (μm): 0.5, temporal filter radius (s): 1, cortex depth (μm): 0.5, Visualization line thickness: 2. The output of ADAPT are the edge velocity and cortical actin density spatiotemporal maps.

The normalized cross-correlation matrices were calculated from the edge velocity and cortical actin density spatiotemporal maps as described in (Machacek et al., 2009; Maeda et al., 2008). The *cross-correlation value c*_(*x, y*)_ at a coordinate (*x, y*) within the cross-correlation map, *c*, of two images *I*_2_ and *I*_2_, is calculated as follows: 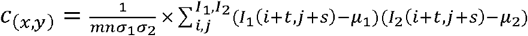, where *m*×*n* are the dimensions of the images *I*_1_ and *I*_2_, and *σ*_2_ are the standard deviations of *I*_1_ and *I*_2_, *μ*_1_ and *μ*_2_ are the means of *I*_1_ and *I*_2_, and *t* = *x* – *m*/2 and *s* = *y* – *n*/2. For the calculation of the autocorrelation, *I*_1_ = *I*_2_. To calculate only the temporal or the spatial cross-correlation coefficient, only a vector of the cross-correlation matrix *C*(*s, t*) was considered: *C*(*0, t*) for the temporal cross-correlation or *C*(*s*, 0) for the spatial cross-correlation.

To calculate the *persistence times and widths*, we performed autocorrelation of the edge velocity map, *C_vel vel_*(*s, t*). By definition, after normalization, *C_vel vel_*(0, 0) = 1. We fitted an exponential to the elements of the autocorrelation matrix *C_vel vel_*(0, *t*) or *C_vel vel_*(*x*, 0). The mean persistence time was defined as 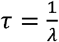 and the mean protrusion width as 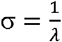 where *λ* are the decay constants of the fitted exponentials. This is similar to the persistence time defined in the study of cell trajectories defined as persistent Brownian motion (Selmeczi et al., 2005). To extract the decay constants, linear fits were performed in log space for the correlation range = (0.95-0.25). This algorithm was implemented in the home-made Python script “ADAPT_processing.ipynb”.

## Supporting information

Supplementary Theory information

## Acknowledgements

We thank Li Wang, Jian Shi, Yan-Jun Liu and Rafaële Attia for help with photolithography and microfabrication; Jérémie Francfort for help and advice with theory and analysis; Damien Cuvelier, Valeria Venturini, Kotryna Vaidžiulytė, Mathieu Coppey, Manos Mavrakis, Larisa Venkova, Ayako Yamada, Rikki Garner and Vahe Galystan for fruitful discussions and advices; Anne-Sophie Mace and Mathieu Maurin for advice on image analysis and technical assistance; and Kevin Gateau for help with 3D-printing prototyping. We thank all the Piel, Ruprech, and Wieser lab members for hosting the project and helping the authors, in particular but not limited to: Matthieu Deygas, Nishit Srivastava, Ido Lavi, Aastha Mathur, Pablo Sáez, Pablo Vargas, Clotilde Cadart, Lucie Barbier, Nico Carpi, Alice Williart, Christian Knapp, Fabio Pezzano, Senda Jiménez Delgado, Merche Rivas Jiménez. We thank the cohort and teachers at the 2019 Physical Biology of the Cell (PBoC) course at EMBL in Woods Hole for input and productive exchanges, made possible by funding from the Boehringer Ingelheim Fonds, the bourses doctorales de mobilité de l’Université de Paris (BDMU), and the Horace W. Stunkard Scholarship Fund. This work was supported by a French Agence Nationale de la Recherche (ANR) grant to MP (ANR-19-CE13-0030). This work has also received the support of Institut Pierre-Gilles de Gennes-IPGG (Equipement d’Excellence, “Investissements d’avenir”, program ANR-10-EQPX-34) and laboratoire d’excellence, “Investissements d’avenir” program ANR-10-IDEX-0001-02 PSL and ANR-10-LABX-31. We also thank the staff members at the Nikon Imaging centre at Institut Curie, the technological platform at Institut Pierre-Gilles de Gennes (IPGG), and the technological platform at Institut de Ciències Fotòniques (ICFO) for providing equipment and technical assistance. JMGA has received funding from Inserm (ITMO Cancer FDV2016), Fondation ARC pour la recherche sur le cancer (DOC20190508743), and a short-Term EMBO fellowship (7873).

## Author contributions

Conceptualization, J.M.G.A. and M.P.; Methodology, J.M.G.A., S.G., G.S., A.L., and C.C; Software, J.M.G.A. and S.G.; Formal Analysis, J.M.G.A. and S.G.; Investigation, J.M.G.A., J.Z., S.G., A.L. and C.J.C.; Resources, J.Z., L.R., D.M., V.R., S.W., R.V. and M.P.; Data Curation, J.M.G.A.; Writing – Original Draft, J.M.G.A., R.V. and M.P.; Writing – Review & Editing, J.M.G.A., R.V. and M.P.; Visualization, J.M.G.A. and S.G.; Supervision, D.M., V.R., S.W., R.V. and M.P.; Project Administration, J.M.G.A. and M.P.; Funding Acquisition, J.M.G.A. and M.P.

---

**Figure S1:**
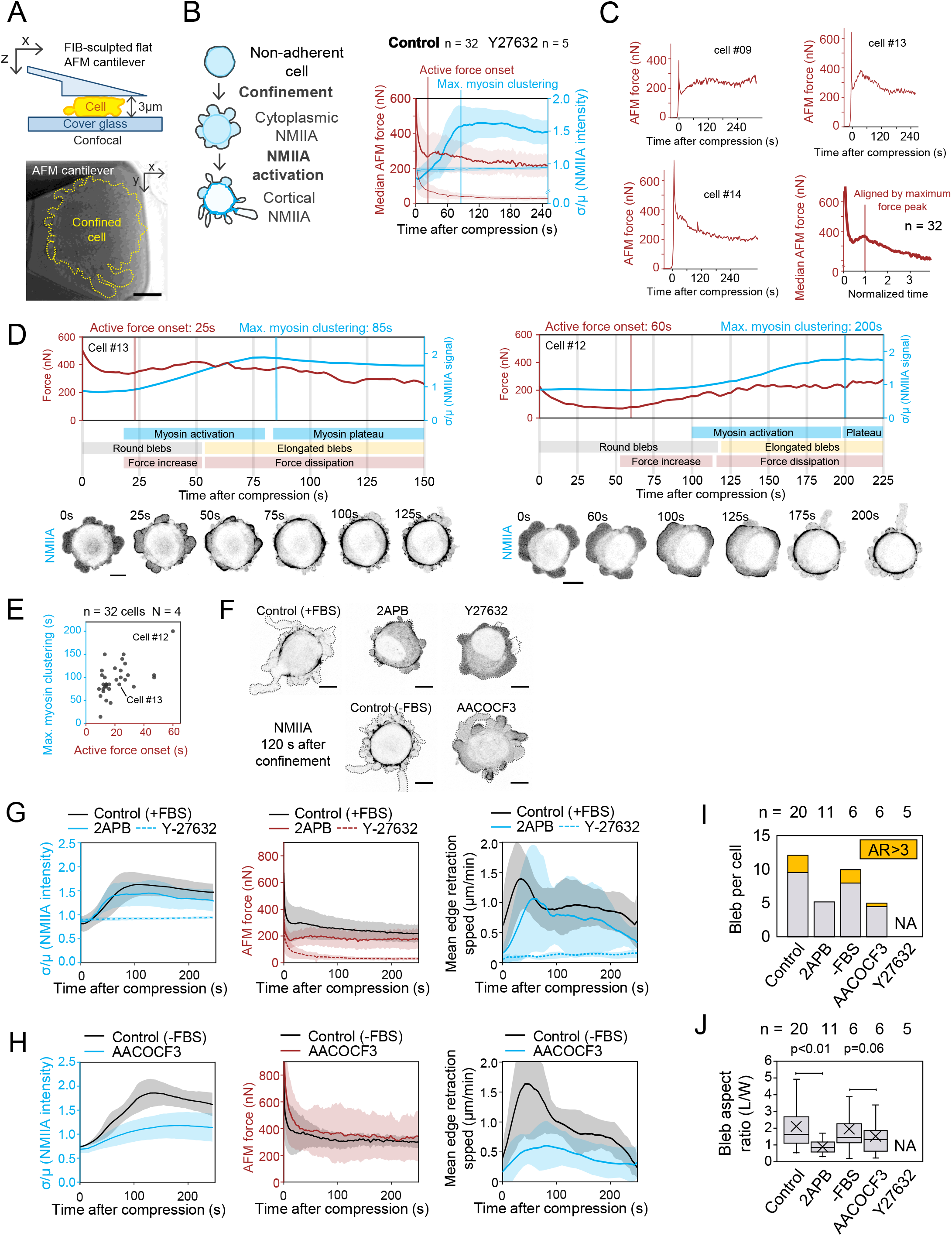
Force and myosin response to AFM compression. Related to Figure 1. **A:** Top, schematic representation of a cell confined to 3μm under a FIB-sculpted flat AFM cantilever combined with confocal microscopy. The measurements of the force exerted by cells on the cantilever and the confocal images are recorded simultaneously. Bottom, phase-contrast image of an AFM-confined HeLa cell. Yellow dashed line, cell outline. Scale bar, 10μm. **B:** Left, Schematic representation of myosin II dynamics upon cell confinement. Non-muscle myosin IIA (NMIIA, cyan) is mostly cytoplasmic in non-adherent HeLa cells. Confinement promotes a transition from cytoplasmic globular NMIIA to cortical phosphorylated myosin minifilaments. This signal clustering can be captured by the coefficient of variation of the background-subtracted NMIIA intensity. Right, AFM force feedback (red, mean ± s.d.) and coefficient of variation of the NMIIA intensity (σ/μ, cyan, mean ± s.d.) for control cells (bold red and blue lines, n = 32) and 10μM Y-27632-treated cells (light red lines, n = 5). Solide lines are mean, shaded area s.d. Active force onset, change of trend of the median AFM force. Maximum myosin clustering, timepoint at which the coefficient of variation of the NMIIA intensity plateaus. **C:** Top and bottom left, three representative AFM force curves (nN) during compression, showing the variability of behaviours observed. Bottom right, median AFM force curve calculated by aligning by the maximum force peak, with normalized time (×1/t_maximum force peak_). **D:** Two single-cell representative examples of the response to confinement. Top, time response of AFM force (red line, nN) and coefficient of variation of the NMIIA intensity (cyan line). Middle, summary of the cell behaviour (cyan, myosin response; gray and yellow, bleb morphology; red, force response). Bottom, fluorescent confocal images 3 μm-confined HeLa *MYH9-eGFP* (NMIIA). Time stamp, time from confinement in seconds. Scale bar, 10 μm. **E:** Time of active force onset (change of trend in the AFM force curve) plotted against the time of maximum myosin clustering time (start of the plateau of the coefficient of variation of the myosin signal) (n=32 cells). **F**: Representative fluorescence confocal images of 3 μm-confined HeLa *MYH9*-eGFP (NMIIA) cells taken 120 s after confinement under different conditions: control, 2-APB 100μM, Y-27632 10μM, control without FBS, and AACOCF3 20μM. Scale bar, 10 μm. **G**: Description of cell response as a function of time from confinement in control + FBS (n = 20), 2-APB 100μM (n = 11), and Y-27632 10μM (n = 5) conditions. Solid lines, average. Shades, standard deviation. Left, σ/μ of NMIIA intensity. Middle, average AFM force (nN). Right, average cell edge retraction speed (μm/min). **H**: Description of cell response as a function of time from confinement in control - FBS (n = 6), and AACOCF3 20μM (n = 6) conditions. Solid lines, average. Shades, standard deviation. Left, σ/μ of NMIIA intensity. Middle, average AFM force (nN). Right, average cell edge retraction speed (μm/min). **I**: Number of blebs per cell during 300s in control, 2-APB 100μM, Y-27632 10μM, control without FBS, and AACOCF3 20μM conditions. Grey, blebs with an aspect ratio smaller than 3. Orange, blebs with an aspect ratio greater than 3. **J**: Quartile distribution and mean (cross) of the bleb aspect ratio in control, 2-APB 100μM, Y-27632 10μM, control without FBS, and AACOCF3 20μM conditions. Outliers not plotted. *P*, Welch’s t-test.

**Figure S2:**
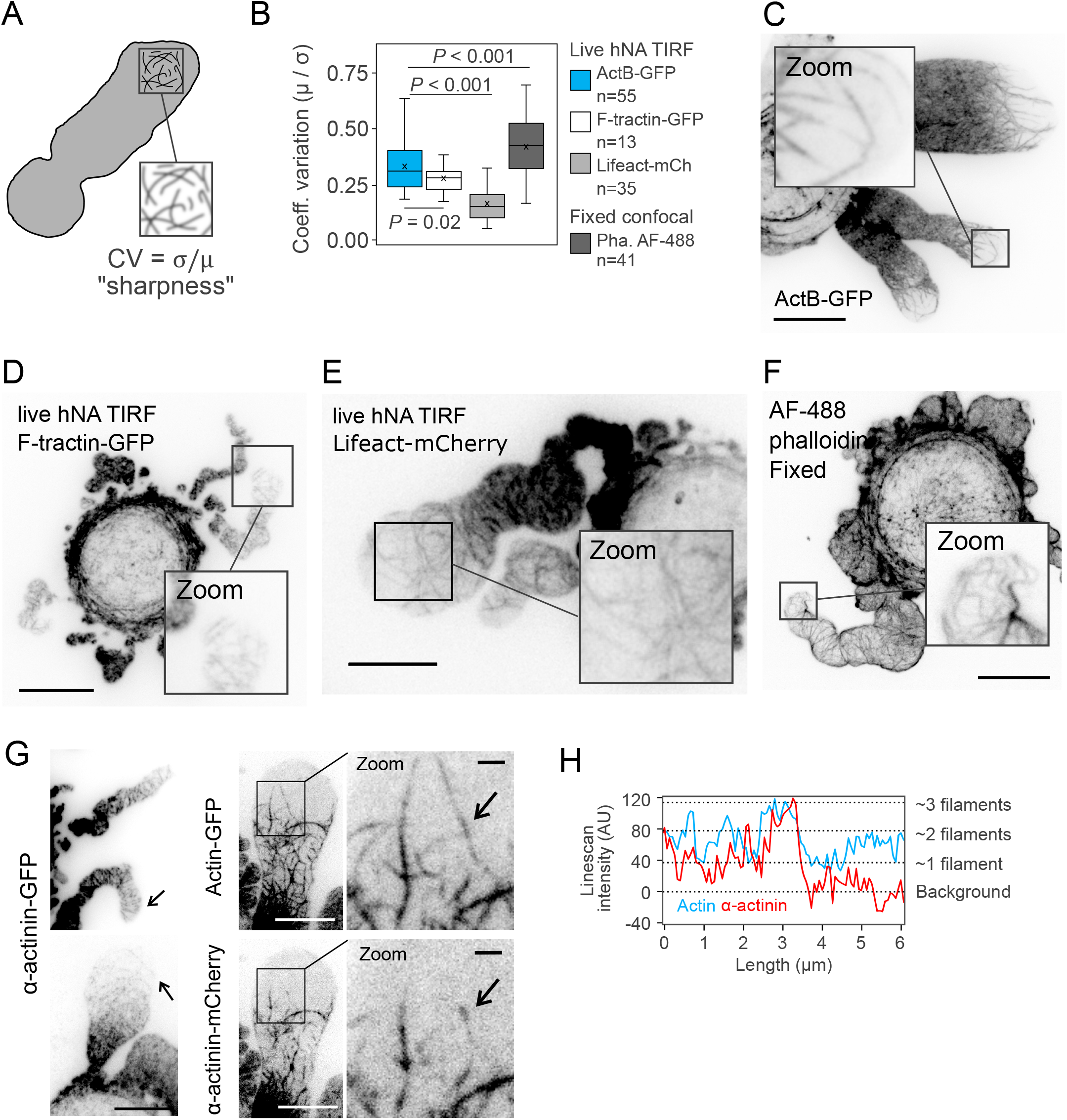
Actin organization in stable blebs with different markers. Related to Figure 2. **A:** Diagram representing the analysis of panel S2B. The coefficient of variation (CV) serves a proxy of the sharpness or focus of the image of the bleb front. **B:** Coefficient of variation of the actin signal measured at the bleb front in ActB-GFP live hNA TIRF (also labelled as actin-GFP, cyan, n = 55 blebs), F-tractin-GFP live hNA TIRF (cyan, n = 13 blebs), lifeact-mCherry live hNA TIRF (cyan, n = 35 blebs), and fixed phalloidin Alexa Fluor 488 imaged on a confocal spinning disk (dark grey, n = 41 blebs). P values, Welch’s T test. **C:** Representative live hNA TIRF image of 3 μm-confined HeLa *ActB*-GFP cells. Bleb square, zoom of the bleb front. Scale bar, 10 μm. **D:** Representative live hNA TIRF image of 3 μm-confined HeLa cells transfected with F-tractin-GFP. Scale bar: 10 μm. **E:** Representative live hNA TIRF image of 3 μm-confined HeLa cells transfected with lifeact-mCherry. Scale bar: 10 μm. **F:** Representative fixed confocal spinning disk image of 3 μm-confined HeLa cells labelled with Alexa Fuor-488 phalloidin. Scale bar: 10 μm. **G:** Left, representative live high numerical aperture TIRF image of 3 μm-confined HeLa cells transfected with alpha-actinin-GFP. Black arrows, filament-like pattern. Scale bar, 10μm. Middle, representative live hNA TIRF image of 3 μm-confined HeLa *ActB*-GFP cells transfected with alpha-actinin-mCherry. Scale bar, 10 μm. Right, zoomed image. Black arrows, sites of filament overlapping and accumulation of alpha-actinin. Scale bar, 1 μm. **H:** Background-subtracted intensity of actin (cyan) and alpha-actinin (red) scanned along the filament pointed in panel S2G. Right, interpretation of the intensity levels.

**Figure S3:**
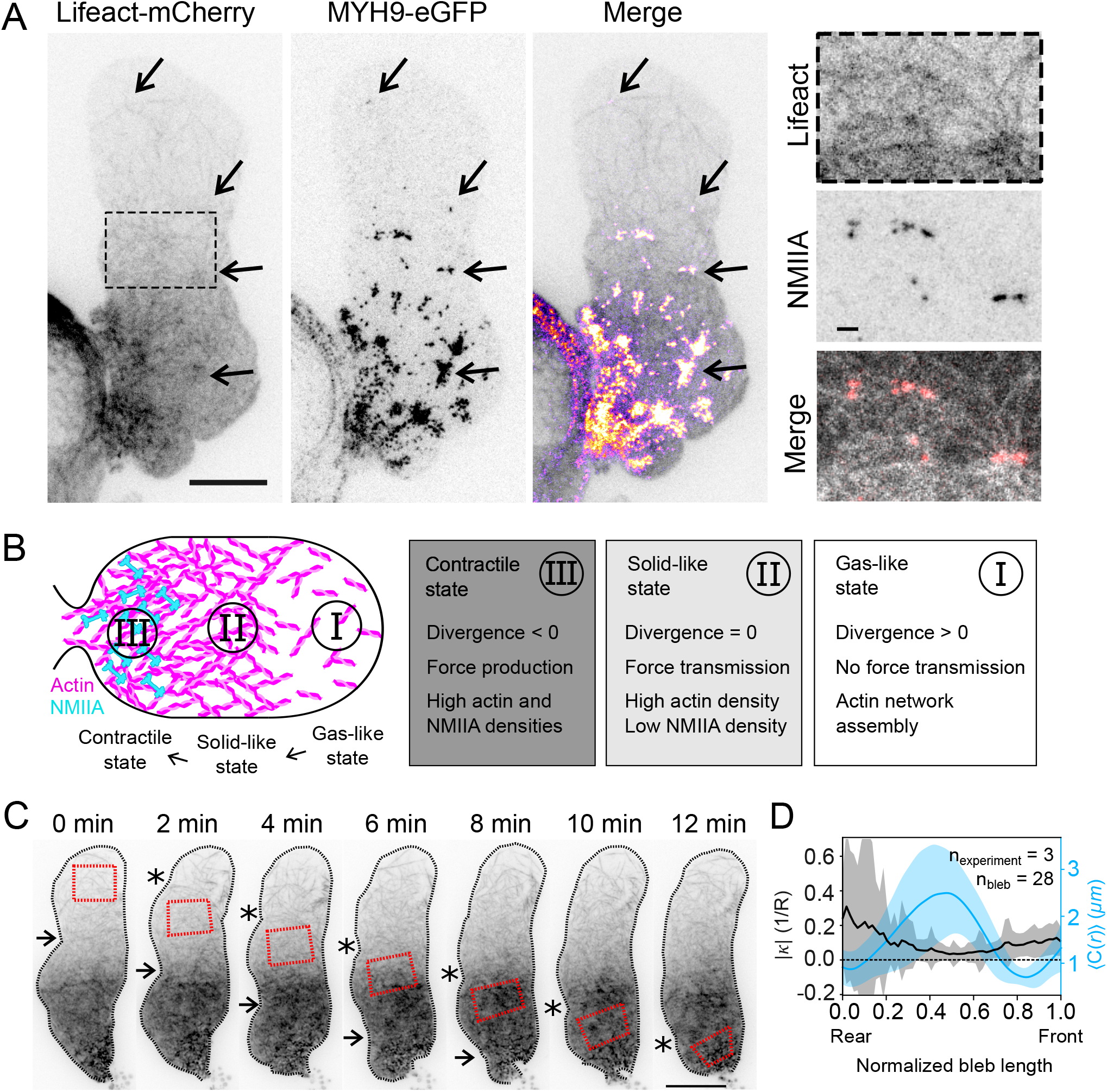
NMIIA molecular assembly in stable blebs. Related to Figure 3. **A:** Left panels, representative hNA TIRF image of 3 μm-confined HeLa lifeact-mCherry (black LUT in merge) *MYH9*-eGFP cells (fire LUT in merge). Arrows, myosin clusters of various sizes. Scale bar, 10μm. Right panels, zoom of the middle, non-contractile bleb region, in lifeact-mCherry (top, Lifeact, black LUT in merge) and *MYH9*-eGFP channels (middle, NMIIA, red LUT in merge). Scale bar, 1 μm. **B:** Schematic representation of the cortical states found spatially organized in stable blebs: ‘gas-like’ front (I), ‘solid-like’ intermediate region (II), and contractile rear (III). Magenta, actin filaments. Cyan, NMIIA, myosin II filaments. **C:** Representative high numerical aperture TIRF time-lapse images of 3 μm-confined HeLa *ActB*-GFP cells. Dashed red polygon, tracking of landmarks (polygon vertices) in the flowing actin network. Dashed back line, bleb contour. Star and arrow, kinks in the bleb contour. Scale bar, 10 μm. **D:** Bleb contour curvature |κ|, 1/radius (μm^-1^, black, mean ± s.d.) and correlation length C(r) (μm, blue, as in panel 3H, mean ± s.d.) along bleb length, averaged over the width and several blebs (N = 3, n_bleb_ = 20).

**Figure S4:**
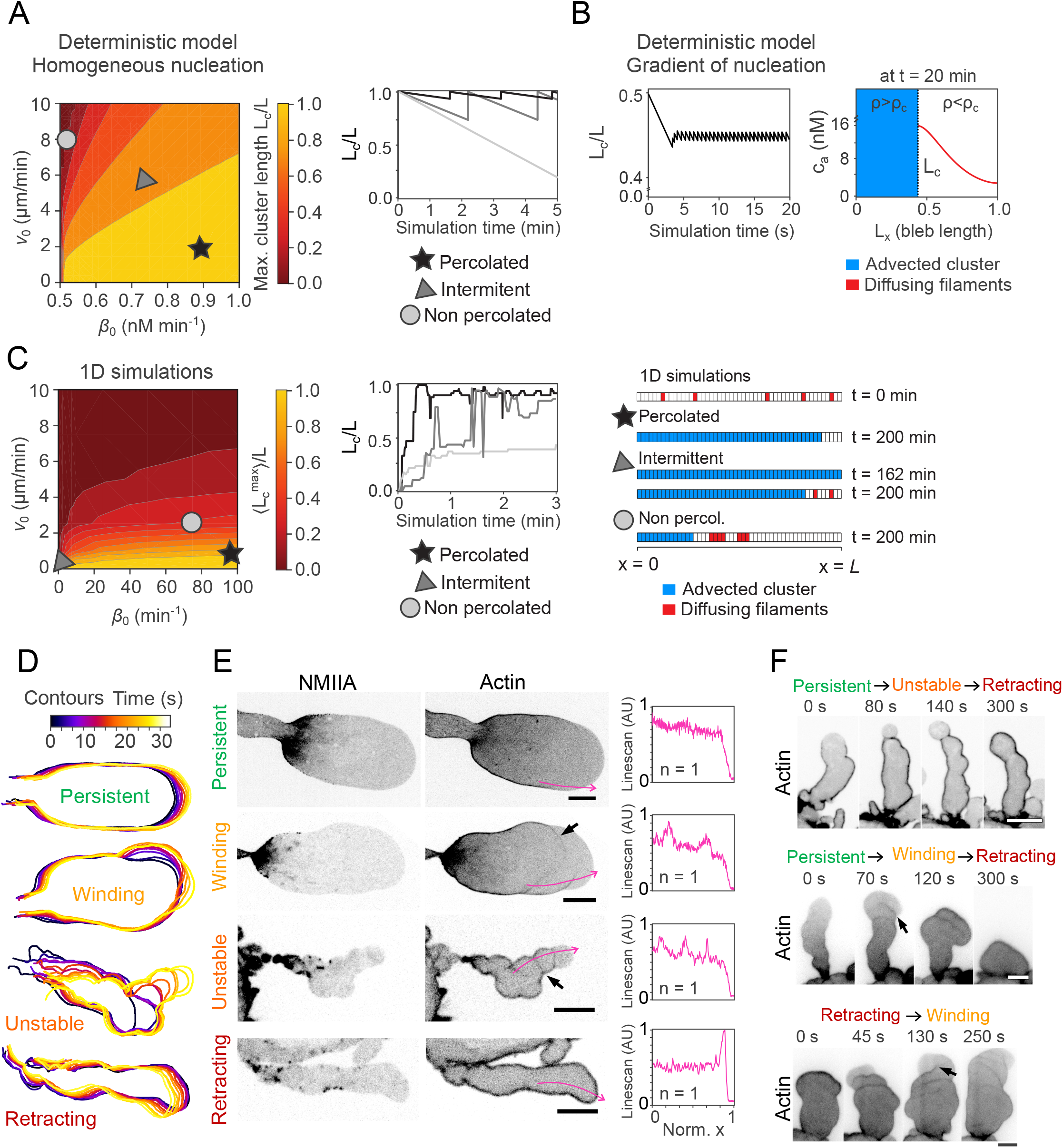
Advected percolation model supplementary figures and bleb regimes. Related to Figure 6. **A:** Outcome of the deterministic advected percolation model with uniform nucleators’ profile. Left, phase diagram of the renormalised maximum cluster length (as shown in the schematics of Figure 6B) varying the advection speed v_0_ and the assembly rate β_0_. Parameters, k_d_ = 0.5 min^1^, D_a_ = 10 μm^2^min^-1^, L_x_ = 50 μm, *c*\* = 1 ⊓M. The details of the numerical integration are reported in the Supplementary Information. Inset, legend. Right, renormalised cluster length, L_c_/L_x_ as a function of elapsed simulation time, for three representative examples pinpointed in the left panel (star, percolated regime; triangle, intermittent regime; circle, non-percolated regime). **B:** Left, renormalised cluster length, L_c_/L as a function of simulation time. Parameters are the same as in panel A, other than β_0_ = 100 *nM/min*, v_0_ = 1 μ*m/min* and *c*\* = 0.98 · β_0_/*k_d_* · *e*^-*L_x_*/2*L_N_*^. Right, concentration profile of actin along the bleb length at t = 20 min. The details of the numerical integration are discussed in the SI. **C:** Left, phase diagram of the stochastic advected percolation model as a function of the advection speed v_0_ and the assembly rate β_0_. Legend is shown as inset. Middle, representative dynamics of the renormalised cluster length for the three cases pinpointed in the left panel. Time, elapsed simulation time. Right, lattice configurations for the three cases plotted in the middle panel. Time, elapsed simulation time. Blue, sites belonging to the cluster. Red, free filaments. White, empty sites. **D:** Colour-coded time projection of contours of four representative persistent, winding, unstable, and retracting blebs from 3 μm-confined HeLa *MYH9*-eGFP Lifeact-mCherry cells. Inset, time-colour correspondence. Representative **E:** Left, fluorescence confocal spinning disk time-lapse images of four representative persistent, winding, unstable, and retracting blebs from HeLa *MYH9*-eGFP (NMIIA, first column) lifeact-mCherry cells (Actin, second column), as shown in panel S4D. Black arrows, sites of transient actin cortex reformation. Magenta arrows, intensity line scans plotted in right panel. Scale bar, 10 μm. Right, normalized actin density line scans from magenta arrows in left panel. Persistent blebs (top row) display a leading edge persistently depleted of actin. Intermittent blebs could display a winding front protruded by membrane tearing waves (with actin cortex scars corresponding to the accumulation of percolated actin filaments at alternating sides) (black arrows on “winding” row). Intermittent blebs could also display a more unstable protrusion if a full cortex was reformed, leaving actin cortex scars across the full bleb width (black arrows on “unstable” row). **F:** Representative fluorescence confocal spinning disk time-lapse images of blebs transitioning between regimes from 3 μm-confined HeLa *ActB*-GFP (top row) or HeLa *MYH9*-eGFP Lifeact-mCherry cells (middle and bottom rows). Time, time elapsed from first frame in seconds. Black arrows, sites of transient actin cortex reformation. Scale bars, 10 μm.

**Figure S5:**
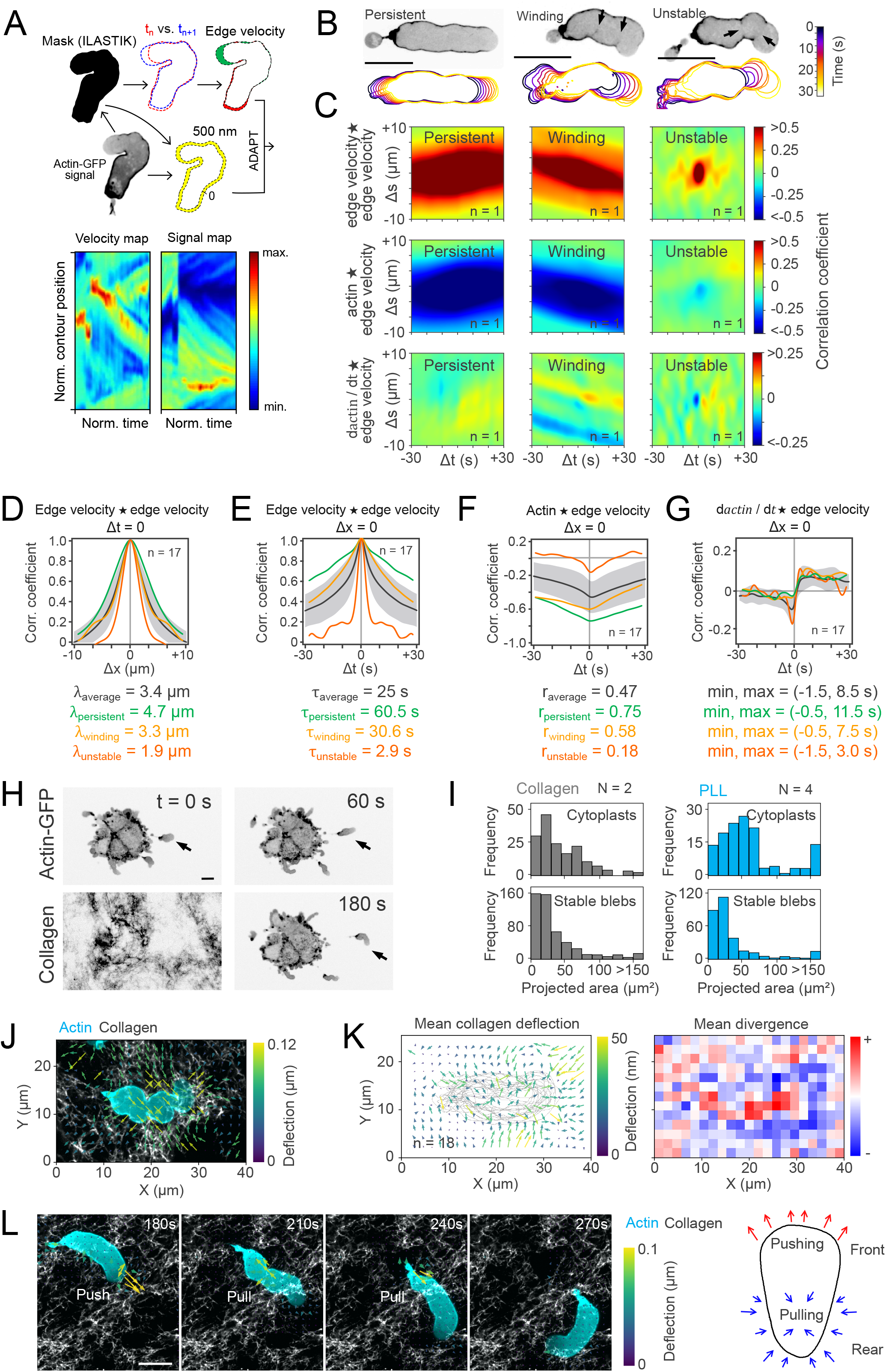
Cytoplast migration in collagen gels. Related to Figure 7. **A:** Pipeline for protrusion and actin cross-correlation analysis of bleb fragments. First, the signal is acquired on actin-GFP. The high background of the actin-GFP signal allows for a good segmentation, which is performed manually for a few time points and then fed into Ilastik to segment the rest of the movie. The Fiji plugin ADAPT (Automated Detection and Analysis of ProTrusions) allows for the analysis of protrusion edge and image segmentation. The actin cortical density is calculated on a 300 nm area beneath the cell contour, on background-subtracted actin-GFP movies. The cross-correlation analysis, normalization, and plotting is performed using a homemade Python algorithm. The output of the pipeline is three cross-correlation matrices: edge velocity ⍰ edge velocity, actin ⍰ edge velocity, and dactin/dt ⍰ edge velocity. **B:** Colour-coded time projection of cytoplast contours and fluorescence confocal spinning disk images of representative migrating cytoplasts from 3μm-confined HeLa *ActB*-GFP cells (Actin-GFP middle plane) under adhesive conditions, displaying persistent (top), winding (middle), and unstable (bottom) migration. Inset, colour code. **C:** Cross-correlation (sliding dot product) matrices of edge velocity ⍰ edge velocity, actin ⍰ edge velocity, and d*actin*/d*t* ⍰ edge velocity for representative examples of persistent, winding, and unstable migration. Inset, colour scale bar, corresponding to correlation coefficient. **D:** Top, edge velocity spatial autocorrelation. Black line, mean ± s.d. (n = 17). Coloured lines show edge velocity autocorrelation curves of three representative persistent (green), winding (yellow), and unstable (orange) cytoplasts. Bottom, decay lengths of the average curve (grey) and three representative persistent (green), winding (yellow), and unstable (orange) cytoplasts. **E:** Top, edge velocity temporal autocorrelation. Black line, mean ± s.d. (n = 17). Coloured lines show edge velocity autocorrelation curves of three representative persistent (green), winding (yellow), and unstable (orange) cytoplasts. Bottom, decay lengths of the average curve (grey) and three representative persistent (green), winding (yellow), and unstable (orange) cytoplasts. **F:** Top, actin ⍰ edge velocity temporal cross-correlation. Black line, mean ± s.d. (n = 17). Coloured lines show cross-correlation curves of three representative persistent (green), winding (yellow), and unstable (orange) cytoplasts. Bottom, values of actin-velocity correlation coefficient of average curve (grey) and three representative persistent (green), winding (yellow), and unstable (orange) cytoplasts. **G:** Top, d*actin/dt* ⍰ edge velocity temporal cross-correlation. Black line, mean ± s.d. (n = 17). Coloured lines show cross-correlation curves of three representative persistent (green), winding (yellow), and unstable (orange) cytoplasts. Bottom, minima and maxima near the vertical axis of the average curve (grey) and three representative persistent (green), winding (yellow), and unstable (orange) cytoplasts. **H:** Representative fluorescence confocal spinning disk time-lapse images of 3μm-confined HeLa *ActB*-GFP (Actin-GFP) cells embedded in a collagen gel (Collagen). Black arrow, fragmenting bleb. Scale bar, 10 μm. **I:** Histogram of projected area of 3μm-confined cytoplasts (top row) or stable blebs (bottom row) embedded in 2mg/ml collagen (grey, left column) or in a PLL-coated chamber (cyan, right column). **J:** Representative fluorescence confocal spinning disk image of a cytoplast from 3μm-confined HeLa *ActB*-GFP (Actin-GFP, cyan) cells embedded in a collagen gel (Collagen, gray), and collagen deflection map, resulting from comparing with PIV the collagen channel during and after cytoplast migration. Arrow colours, deflection magnitude. Inset, colour scale bar (Deflection, μm). Cyan, actin channel. Grey, collagen channel. **K:** Left, mean collagen deflection map as in panel S5J (n = 18). Arrow colours, deflection magnitude. Grey, cytoplast contours. Inset, colour scale bar (Deflection, μm). Right, divergence 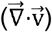 of the mean collagen deflection map shown on the left. Inset, colour scale bar (sign of the divergence). **L:** Left, representative collagen deflection map during fragment migration. Arrow colours, deflection magnitude. Cyan, actin channel. Grey, collagen channel. Right, diagram showing inferred forces applied to the collagen matrix.

**Figure S6:**
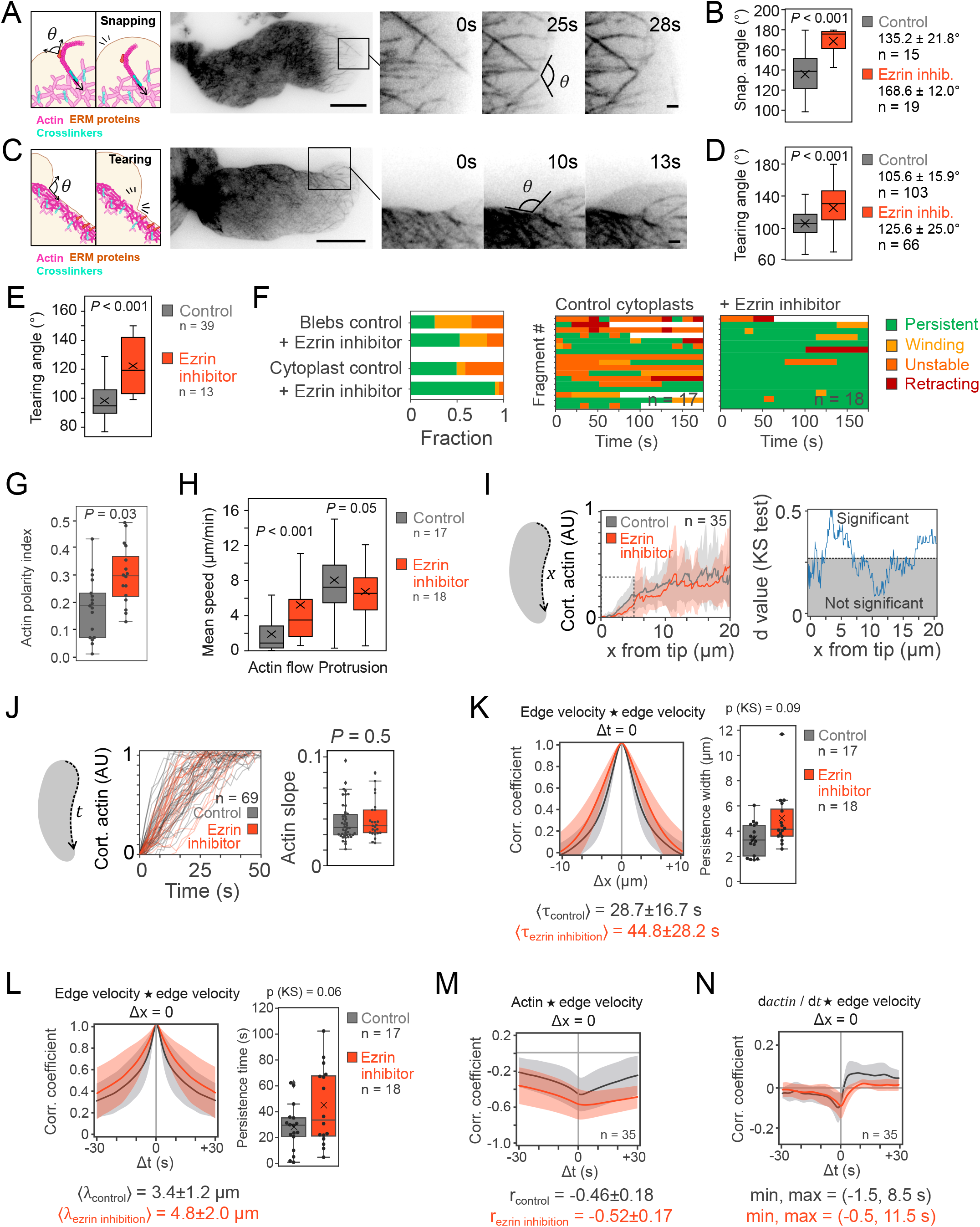
Effect of ezrin inhibition in bleb protrusion and cytoplast migration. Related to Figure 7. **A:** Left, schematic representing the event of membrane snapping. Middle, representative high numerical aperture TIRF images of HeLa *ActB*-GFP cells with an event of membrane deformation and snapping. Scale bar, 10μm. Right, zoomed time-lapse images of membrane deformation and snapping (measured angle marked on the image). Time stamp, time elapsed from first frame in seconds. Scale bar, 1μm. **B:** Bleb membrane snapping angle (°) in control (grey, n = 15) and ezrin inhibitor NSC668994 10μM (orange, n = 19) conditions. P value, Welch’s T test. Cross, average. Bar, median. **C:** Left, schematic representing the event of membrane tearing. Middle, representative high numerical aperture TIRF images of HeLa *ActB*-GFP cells with an event of membrane tearing. Scale bar, 10μm. Right, zoomed time-lapse images of membrane deformation (measured angle marked on the image). Time stamp, time elapsed from first frame in seconds. Scale bar, 1μm. **D:** Bleb membrane tearing angle (°) in control (grey, n = 103) and ezrin inhibitor NSC668994 10μM (orange, n = 66) conditions. P value, Welch’s T test. Cross, average. Bar, median. **E:** Cytoplast membrane tearing angle (°) in control (grey, n = 39) and ezrin inhibitor NSC668994 10μM (orange, n = 13) conditions. P value, Welch’s T test. Cross, average. Bar, median. **F:** Left, time-prevalence of unstable (orange), winding (yellow), persistent (green) phenotypes in control and ezrin inhibitor conditions in blebs (control, n = 176 blebs; NSC668994, n = 68) and cytoplasts (control, n = 17 blebs; NSC668994, n = 18). Right, representation of cytoplast blebbing regimes (retracting, unstable, winding, persistent) in control and ezrin inhibitor NSC668994 10μM conditions over 350 frames. **G:** Average actin polarity in control (grey, n = 17) and ezrin inhibitor NSC668994 10μM (orange, n = 18) conditions. P value, Welch’s T test. Cross, average. Bar, median. **H:** Retrograde cortical flow or protrusion speeds in control (grey, n = 17) and ezrin inhibitor NSC668994 10μM (orange, n = 18) conditions. P value, Welch’s T test. Cross, average. Bar, median. **I:** Left, actin cortical gradient from the bleb tip in control (grey, mean ± SD, n = 17), and ezrin inhibitor NSC668994 at 10μM conditions (orange, mean ± s.d., n = 18). Right, d-value (from Kolmogorov–Smirnov test) comparing control and ezrin inhibitor conditions at every point. White area, significant difference (alpha < 0.05). **J:** Left, cortical actin density in cortical regions flowing from the front to the rear, as a function of time (time=0 is bleb front). Curves are normalized by their plateau. Slope of the curves from normalized density=0.2 to 0.8. P value, Welch’s T test. Right, actin assembly slope (assembly rate). P value, Welch’s T test. Bar, median. **K:** Left, edge velocity spatial autocorrelation function for control (grey, mean ± s.d., n = 17), and ezrin inhibitor NSC668994 at 10μM conditions (orange, mean ± s.d., n = 18). Right, decay lengths of edge velocity spatial autocorrelation. P value, Welch’s T test. Cross, average. Bar, median. **L:** Left, edge velocity temporal autocorrelation function for control (grey, mean ± s.d., n = 17), and ezrin inhibitor NSC668994 at 10μM conditions (orange, mean ± s.d., n = 18). Right, decay lengths of edge velocity temporal autocorrelation. P value, Welch’s T test. Cross, average. Bar, median. **M:** Top, actin ⍰ edge velocity temporal cross-correlation for control (grey, mean ± s.c., n = 17), and ezrin inhibitor NSC668994 at 10μM conditions (orange, mean ± s.d., n = 18). Bottom, actin-velocity Pearson’s correlation coefficient (*r*, mean ± s.d.). **N:** Top, *dactin/dt* ⍰ edge velocity temporal cross-correlation for control (grey, mean ± s.d., n = 17), and ezrin inhibitor NSC668994 at 10μM conditions (orange, mean ± s.d., n = 18). Bottom, minimum and maximum around Δt = 0 (mean ± s.d.).

**Figure S7:**
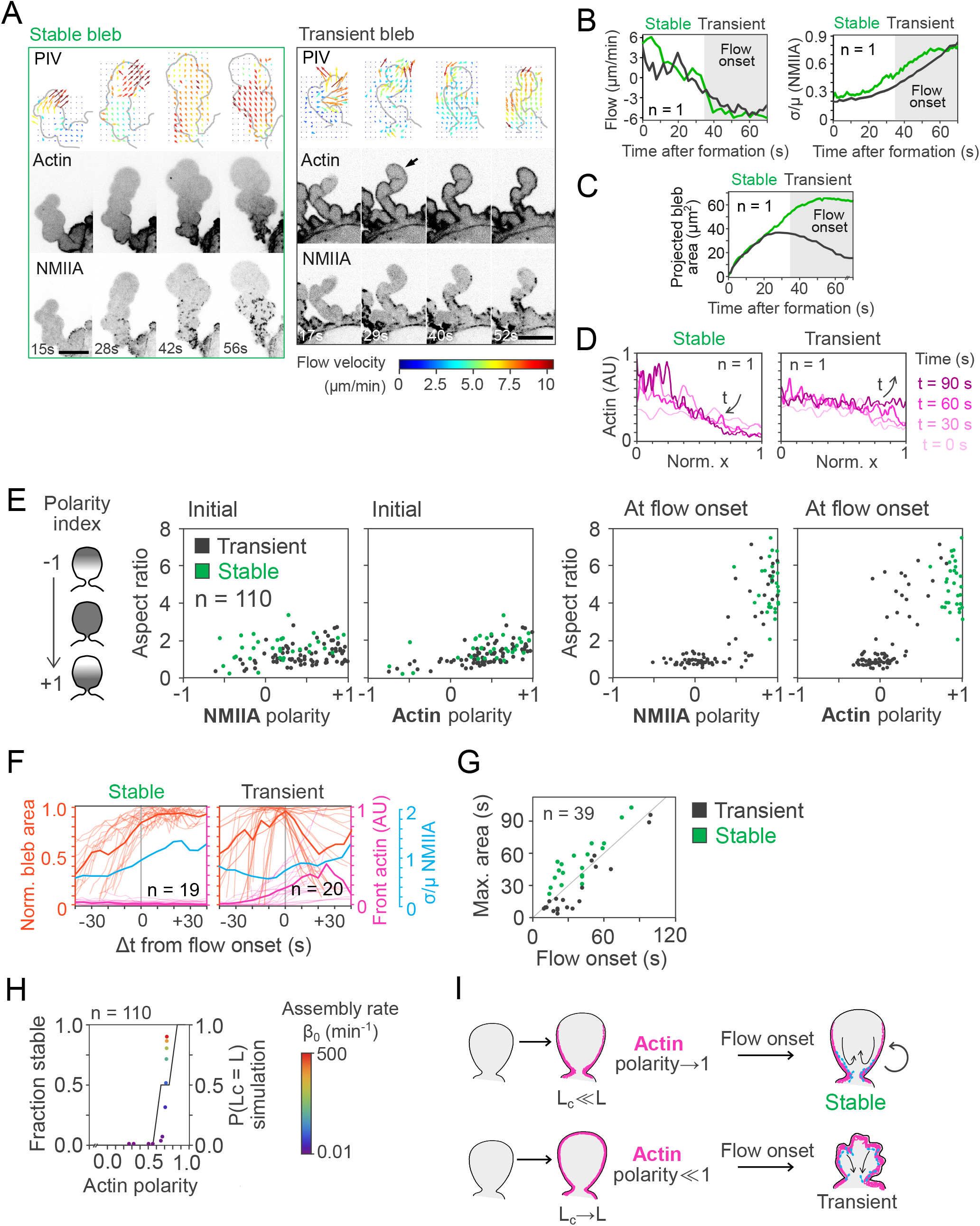
The advected percolation model applied to bleb morphogenesis. Related to Figure 7. **A:** Representative time-lapse images of stable (left panel) or transient bleb formation (right panel) in 3μm-confined HeLa *MYH9*-eGFP Lifeact-mCherry cells. First row (PIV), PIV (particle image velocimetry) applied at the binarized *MYH9*-eGFP (NMIIA) channel. Colours, velocity magnitude, as shown in inset bar. Second row (Actin), fluorescence confocal spinning disk time-lapse images of the Lifeact-mCherry channel. Third row (NMIIA), fluorescence confocal spinning disk time-lapse images of the *MYH9*-eGFP channel. Time stamp, time elapsed from first frame in seconds. Black arrow, cortex formation at the bleb tip. Scale bars, 10 μm. **B:** Average PIV flow speed (left, flow) and coefficient of variation (s.d./mean) of *MYH9-eGFP* signal (right, σ/μ NMIIA) as a function of time elapsed from bleb formation for the stable bleb (green) and transient bleb (grey) shown in panel S7B. Shade, time points after flow onset. **C:** Projected bleb area as a function of time elapsed from bleb formation for the stable bleb (green) and transient bleb (grey) in panel B. Shaded area, time after flow onset. **D:** Actin cortical spatial gradients of the stable (left) and transient (right) blebs shown in panel S7B. X axis, renormalized bleb position (1 corresponding to bleb front). Colour code, time elapsed from bleb formation. **E:** Left, diagram illustrating the polarity index. Accumulation at the bleb base yields a polarity index of +1, whereas homogeneous accumulation yields a polarity index of 0. Middle, initial (1 to 5 s after bleb formation) distribution of bleb aspect ratio (length over width), NMIIA polarity index, and actin polarity index, and coloured by bleb stability (green, stable; gret, transient; n = 110). Right, distribution of bleb aspect ratio (length over width), NMIIA polarity index, and actin polarity index, and coloured by bleb stability at flow onset (green, stable; grey, transient; n = 110). **F:** Normalized bleb area by the maximum (orange),cortical actin intensity at the bleb front at the bleb front (magenta, distal actin), and coefficient of variation (s.d./mean) of *MYH9*-eGFP signal (cyan, σ/μ NMIIA) as a function of time elapsed from flow onset for stable (n = 19, left) and transient blebs (n = 20, right). Bold, average. Shaded lines, individual curves. **G:** Time of membrane stalling as a function of the time of flow onset (n = 39), both measured relative to bleb formation. Colour, bleb stability (grey, transient; green, stable). Line, slope x=y. **H:** Dark line, fraction of stable blebs (green) as a function of actin polarity measured experimentally (n = 110). Coloured dots, actin polarity as a function of the probability of the percolating cluster to reach the bleb length, measured from simulations. Parameters: v_0_=l μm min^-1^, D_a_=25μm^2^ min^-1^ β_0_=(0.01-500) min^-1^. The rest of the parameters are set as in Panel 6B. Inset, colour code for the actin assembly rate *β*_0_ values in the simulations. **I:** Diagram of the proposed model for stable bleb morphogenesis. Contractility and actin assembly compete to determine bleb stability. The onset of contractility depends on the timing of NMIIA contraction (cyan) and actin cortex reformation (magenta). Black line, plasma membrane. If the onset of contractility occurs before the formation of a full cortex (L_c_ « L, actin polarity ≈ 1), the actin will be removed from the front of the bleb continuously, creating a stable gradient and a persistent retrograde flow (Stable, green, top row). If the onset of contractility occurs only after the cortex has completely reformed (L_c_ → L, actin polarity≪1), NMIIA motor activity will retract the bleb (Transient, grey, bottom row).

## Supplementary movies

**Movie S1, corresponding to Fig. 1 and S7F. From transient to stable blebs and to motile bleb fragments. Part 1** (corresponding to Fig 1 A-B): Switch from transient to stable blebs upon confinement; **Part 2** (corresponding to Figure 1C and S7A): Actomyosin dynamics in a transient bleb; **Part 3** (corresponding to Figure 1C and S7A): Actomyosin dynamics in a stable bleb; **Part 4** (corresponding to Fig. 1 D): Formation of motile bleb fragments on PLL-coated glass substrate; **Part 5** (corresponding to Fig. 1F): Examples of motile bleb fragments.

**Movie S2, corresponding to Fig.2. TIRF imaging of actin filaments in confined stable blebs. Part1** (corresponding to Fig 2A): Actin dynamics in a persistent bleb; **Part 2** (corresponding to Fig 2A): Actin dynamics in a winding bleb; **Part 3** (corresponding to Fig. 2B): Actin filament assembly at the bleb tip; **Part 4** (corresponding to Fig. 2E-F): Branching of a single actin filament; **Part 5** (corresponding to Fig. 2F): Collapse of an actin branch; **Part 6** (corresponding to Fig. 2G): bundling of single actin filaments; **Part 7** (corresponding to Fig. S2GH): Actin dynamics with α-actinin staining.

**Movie S3, corresponding to Fig. 3. TIRF imaging of actomyosin dynamics in stable blebs. Part 1** (corresponding to Fig. 3A): Actin and Myosin dynamics in a stable bleb, with PIV and actin segmentation; **Part 2** (corresponding to Fig 3I and S3C): the solid-like state of the actin network shapes the bulk middle part of stable blebs.

**Movie S4, corresponding to Fig. 5**. **Force transmission through the actin network to the front bleb membrane. Part 1** (corresponding to Fig. 5A): actin network deformation; **Part 2** (corresponding to Fig. 5B): actin network rupture; **Part 3** (corresponding to Fig. 5C): membrane snapping upon pulling from an actin filament bundle at the bleb front; **Part 4** (corresponding to Fig. 5D): membrane deformation upon pulling from an actin filament bundle at the bleb front.

**Movie S5, corresponding to Fig. 5G. Perturbation of stable blebs upon inhibition of actin nucleation factors. Part 1** (corresponding to Fig. 5G): Actin filaments dynamics in CK-666 treated stable blebs (Arp2/3 inhibition); **Part 2** (corresponding to Fig. 5G): Actin filaments dynamics in SMIFH2 treated stable blebs (formin inhibition).

**Movie S6, corresponding to Fig. 6, S4 and S5**. **The various dynamics of stable blebs.** (corresponding to Fig. S4E): Actomyosin dynamics in persistent, winding, unstable and retracting blebs.

**Movie S7, corresponding to Fig. 7**. **Motility of bleb fragments in 2D confinement and in collagen gels. Part 1** (corresponding to Fig. 7B): Actin dynamics in a motile bleb fragment migrating in a PLL-coated 2D confined environment; **Part 2** (corresponding to Fig. 7J and S5L): Actin dynamics in motile bleb fragments navigating inside a dense collagen gel; **Part 3** (corresponding to Fig. 7N): detailed actin dynamics in a fragment on High NA TIRF.

There is also a public repository where all the movies corresponding to still images shown in the figures can be downloaded, with names matching with figures and panel numbers.

DOI: 10.6084/m9.figshare.19878253

Address: https://figshare.com/articles/media/Movies/19878253.

