## Supplementary Theory information for "Advected percolation in the actomyosin cortex drives amoeboid cell motility"

### Supplementary Model

#### I. Advected percolation model : mean field description

##### A. General equations and case of uniform concentration of actin nucleators

We formulate an advected percolation model describing the dynamics of actin in blebs. We consider a simplified quasi 1-dimensional geometry where a bleb extends from the cell body ($x = 0$) to its tip ($x = L$), and assume that the concentration of actin fialments $c_a(x,t)$ depends only on the space variable $x$. Following experimental observations, we assume that actin filaments undergo a rigidity percolation transition. This is described at a mean field level by a critical concentration $c^*$ that we assume is given ; for $c_a(x,t) < c^*$ actin filaments are in a dispersed gas-like phase, whereas for $c_a(x,t) \geq c^*$ filaments form a rigid cluster. We will call main actin cluster the set of points such that $c_a(x,t) \geq c^*$ for all $x \in [0, L_c]$. $L_c \leq L$ thus defines the front end of the main actin cluster, which is connected to the bleb rear.

At the front of the bleb ($x \in [L_c, L]$), actin filaments form a dispersed gas-like phase ; we assume that actin filaments can uniformly assemble with assembly rate $\beta_0$, depolymerise with rate $k_d$ and diffuse with diffusion constant $D_a$. The main actin cluster at the bleb rear ($x \in [0, L_c]$) is described simply as a rigid solid. Following experimental observations, we assume that myosin motors, localized at the bleb rear ($x \simeq 0$), impose the rearward advection of the main cluster at constant speed $-v_0$ (with $v_0 > 0$). In principle $v_0$ is set by the balance of contractile forces induced by myosin motors and resistive friction forces of the cluster with its environment ; it is assumed constant hereafter. For simplicity, we neglect actin turnover in the main cluster, which is irrelevant to describe the dynamics of the system, which is quantified by $L_c(t)$ and $c_a(x,t)$ for $x \in [L_c, L]$ ; we thus set as a convention $c_a(x,t) = c^*$ for all $x \in [0, L_c]$.

The spatio-temporal evolution of the concentration of actin filaments $c_a(x,t)$ is therefore given by

$$\begin{cases} \partial_t c_a = 0 \text{ for } x < L_c \\ \partial_t c_a = D_a \Delta c_a + \beta_0 - k_d c_a \text{ for } x \geq L_c \end{cases} . \tag{1}$$

The boundary condition at the bleb front is assumed to be reflective to enforce conservation of actin filaments. At $x = L_c$, we impose flux conservation for both the main cluster and free actin filaments ; we remind that free actin filaments contribute to the main cluster only if $c_a(x =$

$L_c^+, t) = c^*$. This leads to

$$\begin{cases} \partial_x c_a(x,t)|_{x=L} = 0 \\ D_a \partial_x c_a(x,t)|_{x=L_c^+} = (v_0 + \partial_t L_c) c^* = 0 \text{ with } c_a(x = L_c^+, t) < c^* \end{cases} , \tag{2}$$

which fully defines the dynamics for the unknowns $c_a(x,t)$ and $L_c(t)$. In order to solve Eq. 1 we first notice that as long as $c_a(x = L_c^+, t) < c^*$, free filaments are conserved and do not contribute to the main cluster, so that $D\nabla c_a(x = L_c^+, t) = 0$, and therefore $\partial_t L_c = -v_0$. This imposes a moving boundary on the domain $[L_c, L]$ where filaments are in the dispersed phase . In order to solve the equations for $x \in [L_c, L]$, we first redefine the $x$ coordinate as

$$z = \frac{x - L_c}{L - L_c} \tag{3}$$

such that $z \in [0,1]$ and the boundary remains fixed over the dynamics.

By plugging this change of variable into Eq. (1), we obtain

$$\partial_t c_a + \partial_z c_a \partial_t z = \frac{D_a}{(L - L_c)^2} \partial_z^2 c_a + \beta_0 - k_d c_a \,. \tag{4}$$

This can be rewritten as

$$\partial_t c_a + \frac{z-1}{L - L_c} \partial_z c_a \partial_t L_c = \frac{D_a}{(L - L_c)^2} \Delta c_a + \beta_0 - k_d c_a \,, \tag{5}$$

with $\partial_t L_c = -v_0$ as long as $c_a(z = 0, t) < c^*$ ; this fully defines the dynamics of the system, with now fixed boundaries for $z \in [0,1]$.

From these equations we can show that the current model does not admit any acceptable steady state solution with $L_c > 0$ for $\beta_0/k_d > c^*$. Starting from a typical initial solution where the concentration of actin is uniform everywhere and small (typically $c_a(x,0) = c^*/100$ in numerical solutions), the system reaches a limit cycle where $L_c(t)$ exhibits oscillations. As long as $c_a(x = L_c^+, t) < c^*$, the main cluster retracts at constant speed $-v_0$. During this retraction phase, the concentration $c_a(x,t)$ increases everywhere in the domain $[L_c, L]$, provided that $\beta_0/k_d > c^*$. After a finite time $T$, one has $c_a(x = L_c^+, T) = c^*$ ; because $\partial_x c_a(x,T) \geq 0$, at time $T$, one has $c_a(x, T^-) \geq c^*$ everywhere in the domain. The cluster then spans the full bleb and by definition $L_c$ jumps from $L_c(T^-) < L$ to $L_c(T^+) = L$. By definition the length of the cluster is minimal at the end of a retraction phase so that $L_c^{min} = L_c(T^-)$. The cluster next starts retracting again at constant speed, defining a new domain $[L_c, L]$ with dispersed filaments characterized by $c_a(x, T^+) \simeq 0$ for $x \in [L_c, L]$, and the cycle repeats. In fact, it can be shown directly from (5) that uniform solutions with $c_a(x,t)$ independent of $x \in [L_c, L]$ are solutions to the problem with the appropriate boundary conditions. More explicitly, assuming that $L_c(0) = L$, one has the exact solution for $x \in [L_c, L]$:

$$c_a(x,t) = (1 - e^{-k_d t})\beta_0/k_d\,. \tag{6}$$

The period $T$ of oscillations is then given by

$$T = -\frac{1}{k_d}\ln(1 - k_d c^*/\beta_0) \tag{7}$$

and their amplitude by

$$L - L_c^{\min} = v_0 T\,. \tag{8}$$

This exact analysis is confirmed numerically by integration of (5) with Matlab [1] using the Euler algorithm ; while not necessary here because an exact solution is available, this numerical scheme will be needed in the next paragraph. We imposed the kinetic parameters as described in the last paragraph hereby and $c^* = 1\,\mathrm{nM}$ nM, the initial condition $c_a(x,0) = 0.01c^*$ and $L_c(0) = L - 2\Delta x$, where $\Delta x$ is the spatial step. This initial condition corresponds to a cluster filling (almost) the full bleb, and helps reaching faster the steady state. We impose the number of grid points to be $N_{\mathrm{cells}} = 50$ and the time step $\Delta t = 0.01\,\mathrm{min}$.

As the system evolves, we obtained the value of the cluster length $L_c(t)$ and actin concentration profile $c_a(x,t)$ for $x \in [L_c, L]$. Numerical integration was stopped and reset to initial conditions whenever $c_a(x,t) \geq c^*$ for all $x \in [L_c, L]$, as this would correspond to the main cluster filling the full bleb. Note that we restricted our analysis to parameters for which $L_c(t) > 0$ at all times. As expected, the final solution oscillates in time (Fig. S4A right panel). The minimum cluster length $L_c^{\min}$ (end of a retraction phase) is shown in Fig. S4A (left panel) as a function of the advection velocity $v_0$ and assembly rate $\beta_0$, and is consistent with the exact solution (6) .

##### B. Case of polarized concentration of actin nucleators

Importantly, the analysis above predicts, within our hypotheses, that the main cluster always oscillates between $L_c = L_{c,\min}$ and $L_c = L$ and never reaches a steady state with $L_c < L$. This is directly due to our hypothesis of uniform assembly and disassembly of actin filaments. With this hypothesis indeed, during a retraction phase of the cluster, the tip of the cluster ($x \simeq L$) is the first region to be emptied of filaments and left free to assemble new filaments ; because the assembly rate is assumed uniform, the tip region will thus be the region where $c_a(x,t)$ is the largest (in fact the exact solution (6) shows that the profile is flat, because at short time scales diffusion smoothens the profile at the cluster edge). As a consequence, when percolation occurs somewhere because $c_a(x,t) = c^*$, the cluster will always span the full bleb up to $x = L$.

In order to reproduce the variety of actin profiles observed in the experiments, one has thus to relax our hypothesis of uniform assembly rate. We thus introduce nucleators of filament assembly, without presuming their biochemical nature. We start by defining the nucleators concentration over the bleb length as $c_N(x)$. We impose that the total concentration is fixed and equal to $c_N^{tot}$. We assume that the nucleators can diffuse over the bleb length and be advected as well within the cluster. Therefore the dynamical equations are:

$$\begin{cases} \partial_t c_N = D_a \Delta c_N + v_0 \partial_x c_N \text{ if } x \leq L_c \\ \partial_t c_N = D_a \Delta c_N \text{ if } x > L_c \end{cases} . \tag{9}$$

Considering that these processes equilibrate fast compared to the polymerisation/depolymerisation dynamics of actin, we obtain that the steady state profile of nucleators is exponential with typical decay length $L_N = D_a/v_0$ for $x \leq L_c$ and uniform for $x > L_c$. We next assume the assembly rate of filaments to be proportional to the nucleators concentration. To avoid having a discontinuous profile at $x = L_c$, we simply set $\beta(x) = \beta_0 e^{-\frac{x}{L_N}}$. The equation for actin filaments concentration in $x \in [L_c, L]$ now reads:

$$\partial_t c_a = D_a \Delta c_a + \beta_0 e^{-\frac{x}{L_N}} - k_d c_a. \tag{10}$$

The boundary conditions are assumed to be the same as in Eq. 2.

We performed the same change of coordinates as in the case with uniform nucleators profile and we studied the dynamics of the system with the same numerical integration technique as in the previous case. We imposed that whenever the concentration profile reaches $c_a(x = L_c^+, T^-) = c^*$ at a given time $T^-$ (which defines $L_{c,\min} = L_c(T^-)$, the system is reinitialised to $L_c(T^+) = L_{c,\max}$, where $L_{c,\max} = \max\{x,\ c_a(x,T^-) \geq c^*\}$, and $c_a(x,T^+) = 0.97c^*$ for $x \in [L_c(T^+), L_c(T^+) + \Delta x]$, and $c_a(x,T^+) = c_a(x,T^-)$ for $x \in [L_c(T^+) + \Delta x, L]$ ; this last hypothesis is introduced to regularize the numerical scheme. An example of dynamics is shown in Figure S4B, where parameters are chosen as in the model with uniform assembly rate, other than $c^* = 0.98\beta_0/k_d e^{-\frac{L}{2L_N}}$ and $\Delta t = 2\Delta x^2 D_a$. Importantly, the oscillations of $L_c(t)$ are now found to be limited in amplitude because of the gradient of nucleators ; the amplitude increases with $v_0$ and decreases with $L_N$. The maximal length of the main cluster $L_{c,\max}$ satisfies $L_{c,\max} \sim L_N \ln(k_d c^*/\beta_0)$ and is thus critically controlled by actin assembly/disassembly.

#### II. Advected percolation model : stochastic agent based description

To study now a more realistic system and the effects of fluctuations induced by the underlying stochastic kinetics of both diffusion of filaments and filaments assembly/disassembly, we formulated a stochastic version of the advected percolation model. To start, we defined a 1D grid of $N_{cells}+1$ compartments of total length L=50 $\mu$m corresponding to the average bleb neck-tip distance. Each compartment has length $\Delta x = \frac{L}{N_{cells}}$.

In each compartment $i$ we define the number $n_a^i$ of actin filaments $A_i$. Actin filaments can undergo the following reactions: i) assembly with rate $\beta_i = \beta_0 e^{-i/L_N}$, where $\beta_0$ is the basal assembly rate and $L_N$ is the decay length of the nucleators' profile previously defined and ii) degradation with rate $k_d$ such that:

$$\emptyset \xrightarrow{\beta} A_i \tag{11}$$

$$A_i \xrightarrow{k_d} \emptyset \tag{12}$$

We assume that actin filaments can form clusters, indexed by $\alpha$, and defined by the adjacent sites with an actin occupation number different than zero. Single actin filaments can also diffuse between compartments and this can be modelled as $2N_{cells}$ reactions such that:

$$A_{i-1} \underset{d}{\overset{d}{\rightleftharpoons}} A_i \underset{d}{\overset{d}{\rightleftharpoons}} A_{i+1} \tag{13}$$

where $i$ is the compartment index and $d$ is the diffusion rate which is given by $D = \frac{D_a}{\Delta x^2}$, where $D_a$ is the diffusion constant. We define a cluster as the set of filaments occupying neighboring sites ; by definition a cluster is rigid and we assume that each cluster diffuses as a whole, with:

$$D_a^\alpha = \frac{D_a}{l^\alpha \Delta x^2}, \tag{14}$$

where $\alpha$ is the cluster index and $l^\alpha$ is the total length of the $\alpha - th$ cluster. The main cluster is by definition the cluster to which belongs the site $i = 1$ ; this main cluster does not perform diffusion but is advected at constant rate $\nu = \frac{v_0}{\Delta x}$. We finally impose reflective boundary conditions on the right boundary $x = L$.

To reproduce the chemical kinetics, we implemented the Gillespie algorithm [2] which works as follows:

1. once all parameters are set, initialise the system to $N_0 = 5$ ($N_0 = 50$ in the 2D case instead) molecules randomly distributed over the domain;

2. compute one step of the Gillespie algorithm, described in detail in the next paragraph;

3. identify the clusters and their features (i.e., length, first and last compartment, total number of filaments) via the built-in Matlab library `bwlabel()` [1] and rescale the diffusion constants for each compartment according to the clusters size;

4. go back to point 2 and continue until a time $t_{max}$ is reached.

##### A. Gillespie algorithm

To implement the Gillespie algorithm, one has to compute the propensities of each reaction. In our model, the propensities for each compartment $i$ are defined as:

- assembly for each compartment: $a_1^i = \beta^i = \beta_0 e^{-i/L_N}$;

- degradation for each compartment: $a_2^i = k_d n_a^i$;

- cluster diffusion on the left and on the right for each cluster: $a_3^\alpha = a_4^\alpha = D_a^\alpha n_a^\alpha$;

- advection for the first compartment only: $a_5 = \frac{v_0}{\Delta x}$.

We then compute the total propensity as:

$$a_0 = \sum_i (a_1^i + a_2^i + a_3^i + a_4^i) + a_5. \tag{15}$$

We generate two random numbers uniformly distributed in $I = [0, 1]$, $r_1$ and $r_2$, s.t.:

- $r_1$ defines the time interval in which the reaction takes place, $\tau = -\frac{1}{a_0} \log r_1$;

- $r_2$ selects the reaction $\mu$ taking place in this time interval s.t. $\mu$ is the first integer satisfying $r_2 < \frac{\sum_{k=1}^{\mu} a_k}{a_0}$ (where $k$ includes both the superscripts and subscripts of the propensities);

- we modify the number of molecules in the selected compartment according to the chosen reaction i.e.: i) creation, $n_a^i \to n_a^i + 1$; ii) degradation, $n_a^i \to n_a^i - 1$; iii) cluster diffusion on the right/left by translating the entire cluster in space of one site on the right/left hand-side, except for the last one, given the reflective boundary conditon; iv) advection for the first compartment, $n_a^1 \to n_a^1 - 1$;

- we update the time as $t' = t + \tau$ and start over again.

We store the time correlation between the number of particles at time $t$ and the initial condition, i.e.:

$$\mathcal{C}(t,0) = \langle \underline{n}(t) \cdot \underline{n}(0) \rangle - \langle \underline{n}(t) \rangle \langle \underline{n}(0) \rangle, \tag{16}$$

where the average $\langle \cdot \rangle$ is taken over the lattice. We monitor the average over time of the correlation function and start storing the trajectories of the length of the cluster attached to the left hand-side of the lattice, $L_c$, as $\langle \mathcal{C}(t,0) \rangle_t < \epsilon = 0.05$. We repeated this algorithm for different values of the advection velocity, $v_0$, and the amplitude of the assembly rate, $\beta_0$, and reported the results in Fig. S4C along with three time evolutions for the cluster length, $L_c$, and the typical lattice configurations at different interesting time points.

We used the same rationale for the 2D simulations by defining a 2D rectangular lattice whose sides are $L_x = 50$ $\mu$m and $L_y = 10$ $\mu$m and made of $(N_x + 1) \times (N_y + 1)$ sites of length $\Delta x$ and $\Delta y$ respectively. All reactions are defined the same as in the one-dimensional case, except for the diffusion, which can now take place in the four directions, i.e., left, right, up and down, with reflective boundary conditions on all sides other than the left hand-side. Also, the diffusion constant is now rescaled by the cluster area, $\mathcal{A}_\alpha$.

The clusters are defined the same as in the one-dimensional case. In the two-dimensional case as well, we monitored the decay of the time-correlation function of the number of molecules in the system and then stored the largest x-coordinate among the sites belonging the cluster attached to the left boundary, $L_c^{max}$, the smallest x-coordinate among the sites belonging the cluster attached to the left boundary, $L_c^{min}$, and the average x-position of the rightmost contour of the cluster attached to the left boundary, $\langle L_c \rangle$. The results are shown in the Main Text.

#### III. Main parameters

We fixed the parameters in our models by using the experimental measurements reported in the Main Text for $L_x = 50$ $\mu$m, $L_y = 10$ $\mu$m and $D_a = 10$ $\mu$m$^2$/min. We set the depolymerisation rate $k_d = 0.5$ min$^{-1}$, the same order of magnitude as the lowest value reported in literature for the actin cortex in non-migrating cells [3]. In our simulations we varied the advection speed $v_0$ in the same range as the experimental observed values and the intensity of the assembly rate $\beta_0$, which is not accessible experimentally to our knowledge, in a range where all different scenarios could be recovered. For the deterministic model instead we assumed $c^* = 1$ nM in the case with uniform assembly rate and $c^* = 0.98e^{-L_x/2L_N}$ in the case with non-uniform assembly rate.

[1] MATLAB, (R2018b). Natick, Massachusetts: The MathWorks Inc., 2019.

[2] D. T. Gillespie, "Exact stochastic simulation of coupled chemical reactions," The journal of physical chemistry, vol. 81, no. 25, pp. 2340–2361, 1977.

[3] M. Fritzsche, A. Lewalle, T. Duke, K. Kruse, and G. Charras, "Analysis of turnover dynamics of the submembranous actin cortex," Molecular biology of the cell, vol. 24, no. 6, pp. 757–767, 2013.
